# Prediction of Conformational Ensembles and Structural Effects of State-Switching Allosteric Mutants in the Protein Kinases Using Comparative Analysis of AlphaFold2 Adaptations with Sequence Masking and Shallow Subsampling

**DOI:** 10.1101/2024.05.17.594786

**Authors:** Nishank Raisinghani, Mohammed Alshahrani, Grace Gupta, Hao Tian, Sian Xiao, Peng Tao, Gennady Verkhivker

## Abstract

Despite the success of AlphaFold2 approaches in predicting single protein structures, these methods showed intrinsic limitations in predicting multiple functional conformations of allosteric proteins and have been challenged to accurately capture of the effects of single point mutations that induced significant structural changes. We systematically examined several implementations of AlphaFold2 methods to predict conformational ensembles for state-switching mutants of the ABL kinase. The results revealed that a combination of randomized alanine sequence masking with shallow multiple sequence alignment subsampling can significantly expand the conformational diversity of the predicted structural ensembles and capture shifts in populations of the active and inactive ABL states. Consistent with the NMR experiments, the predicted conformational ensembles for M309L/L320I and M309L/H415P ABL mutants that perturb the regulatory spine networks featured the increased population of the fully closed inactive state. On the other hand, the predicted conformational ensembles for the G269E/M309L/T334I and M309L/L320I/T334I triple ABL mutants that share activating T334I gate-keeper substitution are dominated by the active ABL form. The proposed adaptation of AlphaFold can reproduce the experimentally observed mutation-induced redistributions in the relative populations of the active and inactive ABL states and capture the effects of regulatory mutations on allosteric structural rearrangements of the kinase domain. The ensemble-based network analysis complemented AlphaFold predictions by revealing allosteric mediating centers that often directly correspond to state-switching mutational sites or reside in their immediate local structural proximity, which may explain the global effect of regulatory mutations on structural changes between the ABL states. This study suggested that attention-based learning of long-range dependencies between sequence positions in homologous folds and deciphering patterns of allosteric interactions may further augment the predictive abilities of AlphaFold methods for modeling of alternative protein sates, conformational ensembles and mutation-induced structural transformations.

## Introduction

The remarkable advancements of AlphaFold2 (AF2) technology in protein structure modeling have heralded a transformative era in structural biology.^1,2^ AF2 leverages evolutionary insights through Multiple Sequence Alignments (MSAs) derived from related protein sequences, and critically employs hierarchical transformer architecture with self-attention mechanisms that enable the discernment of long-range dependencies and interactions within protein sequences.^1,2^ Self-supervised deep learning models, inspired by natural language processing (NLP) architectures, particularly those incorporating attention-based and transformer mechanisms, have emerged as powerful tools for capturing contextual spatial relationships through training on extensive sets of protein sequences.^3,4^ One of the latest breakthroughs in AI-based protein structure predictions is the development of ESMFold, an end-to-end family of transformer protein language models known for their high accuracy in predicting atomic-level protein structures directly from individual protein sequences eliminating the reliance on MSA information.^5^ OmegaFold represents another related approach, utilizing a hybrid model that combines a protein language model with a geometry-guided transformer model.^6^ Despite the impressive accomplishments of the AF2-based methods and self-supervised protein language models in forecasting static protein structures, they have notable constraints when it comes to their applicability and universality in characterizing conformational dynamics, functional protein ensembles, conformational alterations, and allosteric states.^7^ Numerous recent studies have indicated that while AF2 methods excel in predicting individual protein structures, extending this proficiency to accurately forecast conformational ensembles and map allosteric landscapes remains a significant hurdle.^8–12^ The limitations of AF2 methods in predicting multiple protein conformations may stem from an inherent training bias towards experimentally determined, thermodynamically stable structures, and MSAs containing evolutionary information used to infer the ground protein states. This bias may hinder the ability to accurately capture the full range of conformational dynamics and some studies indicated that protein structure prediction capabilities of the AF2 methods are not trivially expandable for characterization of functional conformational ensembles and accurate mapping of allosteric landscapes.^13–17^ Although AF2 methods showed robust performance in predicting the experimentally determined ground conformation for 98 fold switching proteins, they typically failed to detect alternative structures suggesting the inherent AF2 network bias for the most probable conformer rather than an ensemble of relevant functional states.^14^

Several recent adaptations of the AF2 framework for predicting alternative conformational states of proteins involve reducing the depth of the MSA where only a subset of sequences is sampled, resulting in a shallower MSA.^18^ The rationale behind this approach is to enhance the diversity of sequences used for modeling protein structures, potentially capturing a broader range of alternative conformational states of proteins. The SPEACH_AF (Sampling Protein Ensembles and Conformational Heterogeneity with AlphaFold2) approach involves in silico alanine mutagenesis within the MSAs that can broaden the attention network mechanism within AF2 enabling exploration of distinct patterns of coevolved residues associated with alternative conformations.^19^ AF-Cluster method is another adaptation of the AF2 and involves MSA subsampling followed by subsequent clustering of evolutionarily related or functionally similar sequences. This technique enables predictions of alternative protein states and has shown promise in identifying previously unknown fold-switched states that were ultimately validated by the NMR analysis.^20^ However, AF-cluster can predict fold switching from single sequences by associating structures in its training set with homologous sequences rather than discerning coevolutionary patterns unique for alternative states^21^ which could arguably limit predictive power of this approach in modeling functional conformations for state-switching proteins that are absent in the training set.^21^ By employing SPEACH_AF and AF-Cluster implementations of AF2, conformational ensembles of 93 fold-switching proteins with the experimentally determined structures were generated producing over 280,000 models.^22^ Despite extensive sampling coverage, AF2 predicted success was surprisingly modest of 25%, pointing to limitations in effective characterization of functional conformations for the majority of fold switching proteins. Additionally, when >159,000 models were generated for seven fold-switching proteins outside of the training set, AF2 accurately predicted fold switching in only one of the seven targets with moderate confidence.^22^ The development of different AF2 pipelines highlights a notable concentration on adaptations geared towards widening the spectrum of available protein conformations by integrating sequence and evolutionary content discerned from MSAs and structural information obtained from templates which was proven to be particularly successful in applications to protein kinase an GPCR proteins^23^ Recent analysis of AF2 predictions and direct comparison with the experimental crystallographic maps showed that even high-confidence predictions could often differ from the experimental maps indicating that AF2-based computational predictions must be carefully analyzed with experimental structural validation and analysis.^24^ Collectively, recent developments in the field revealed that critical bottlenecks of the AF2 methods and emerging adaptations are to accurately capture functional conformational ensembles, allosteric states and the effects of mutations on both local and global conformational changes in proteins.^25^

Protein kinases are regulatory machines characterized by the multiple functional states of the catalytic domain^26–30^ and present a class of regulatory switches that operate through dynamic equilibrium between the active and inactive states. The abundance of structural data available for protein kinases, mutant structures and their complexes^31,32^ in a variety of functional forms offers unparalleled opportunities for validating and testing AF2 methods. Quantitative characterization of kinase conformational dynamics has been challenging and biophysical approaches such as nanopore tweezers were employed to characterize the conformational dynamics of ABL kinase domain, revealing interconversion between two major conformational states where one conformation comprises three sub-states.^33^ A pioneering method of high-resolution structure determination of low-populated states by coupling NMR spectroscopy-derived pseudo-contact shifts with Carr-Purcell-Meiboom-Gill (CPMG) relaxation dispersion can characterize kinetics and thermodynamics of conformational changes between ground and excited states, leading to accurate structure determination of high-energy structures for adenylate kinase, calmodulin and Src kinase, we find that high-energy PCSs accurately determine high-energy structures.^34^

An impressive array of various simulation approaches was combined with Markov state models (MSM) to explore in details the free energy landscape of ABL kinase beyond conformations that are known from X-ray crystallography.^35–37^ These illuminating studies by Roux and colleagues revealed that long molecular dynamics (MD) simulations and rigorous kinetic analysis could detect hidden meta-stable conformations accessible to tyrosine kinases, showing that the characterization of the conformational landscapes of protein kinases can provide a valid information about possible target conformations for structure-based design. Recent unbiased MD simulations of the ABL-Imatinib binding captured conformational change between the active and inactive states, showing that binding involved four primary conformational states that differ in structural arrangements of the key functional regions and that fully inactive ABL form can be formed and become stable only in the Imatinib-bound kinase.^38^ These pioneering computational studies also highlighted the intrinsic difficulties and limitations of conventional biophysical simulations to accurately characterize transient states and describe kinetics of conformational changes as even extremely long MD simulations coupled with the enhanced sampling approaches often fail to detect functionally relevant conformations of protein kinases and unable to accurately map complex allosteric landscapes underlying kinase regulation and activation.

In general, despite significant structural wealth of the protein kinases the experimental atomistic characterizations of the functionally relevant transient states has been lacking until recently due to large conformational transformations and short-lived kinase intermediates involved in kinetics of the allosteric shifts. The recent NMR studies provided atomistic view of the energy landscape underlying allosteric regulation of ABL kinase showing how structural elements synergistically generate a multilayered allosteric mechanism that enables ABL kinase to switch between distinct allosteric states.^39^ Another groundbreaking study from the same lab employed NMR chemical exchange saturation transfer (CEST) experiments to determine structure of the unbound ABL kinase in the active state and enable structural characterization of two short-lived hidden inactive conformations I_1_ and I_2_ that are intrinsic for the kinase domain and are different from each other in the critical functional regions.^40^ This study showed that Abl switches between the active and two distinct inactive states where the conformational ensemble is exploited by mutants, ligands, post-translational modifications, and inhibitors to regulate the kinase activity and function.^40^ Atomistic MD simulations and MSM kinetic modeling characterized the dynamics of structural changes between these ABL conformational states further highlighting the inherent limitations of biophysical simulations to directly observe transitions to low-populated functional protein states.^41^

Several recent investigations explored the potential of AF2 methodologies for predicting conformational states in protein kinases. AF2-based modeling of 437 human protein kinases in the active form using shallow MSAs of orthologs and close homologs of the query protein showed the robustness of AF2 methods as selected models for each kinase based on the prediction confidence scores of the activation loop residues conformed closely to the substrate-bound experimental structures.^42^ The ability of AF2 methods to predict kinase structures in different conformations at various MSA depths was examined, demonstrating that using lower MSA depths allows for more efficient exploration of alternative kinase conformations.^43^ By exploring different adaptations of the MSA subsampling architecture, a latest insightful study systematically examined the ability of the AF2 method to characterize the conformational distributions of the ABL kinase domain and predict the effects of mutations that decrease the population of the ground state (M290L, L301I, F382V, M290L+L301I), and several mutations that can increase it (E255V, T315I, F382L, E255V+T315I).^44^ This study revealed potentials and also pointed to significant limitations of the AF2 subsampling methods for predicting the effect of mutations on shifting equilibrium between the active and inactive states, showing that the predicted effects of the mutations on the ground state population are often opposite of those seen in experiments. Furthermore, the statistical significance of the results is markedly reduced for the mutations that are known to reduce ground-state populations.^44^ We recently proposed a new AF2 adaptation in which randomized alanine sequence scanning of the entire protein sequence or specific functional regions is combined with the MSA subsampling enabling interpretable atomistic predictions and adequate characterization of the ABL conformational ensembles for the active and inactive states.^45^ These studies suggested that key challenges of the emerging AF2 adaptations are associated with accurate predictions of functional conformations and relative populations of distinct allosteric states rather than simply increasing the conformational diversity of the predicted ensembles.

In the current study, we further develop and expand this AF2 adaptation by exploring targeted alanine masking of the ABL sequence space in functional regions critical for conformational changes between ABL states. Using AF2 approach with MSA subsampling and the AF2 adaptation with random alanine sequence scanning, we perform comparative analysis of these two AF2 implementations in predicting structures, conformational ensembles and populations for a series of functional double and triple ABL mutants that are known to exert distinct functional effects by shifting the equilibrium between the thermodynamically dominant active ABL form and the inactive I_1_ and I_2_ states. The results of our analysis show that AF2 with shallow MSA subsampling can characterize conformational heterogeneity of the ABL active form but generally falls short of predicting mutation-induced structural changes resulting in the inactive ABL states. In some contrast, a combination of targeted alanine sequence masking scanning with shallow MSA subsampling can significantly expand the conformational diversity of the predicted structural ensembles and capture populations of both active and inactive ABL states. In particular, we show that the proposed AF2 enhancement can capture the effects of ABL mutational variants M309L/L320I, M309L/H415P that shift the equilibrium away from the active form and increase population of otherwise hidden I_2_ state. The results reveal that the predicted ensembles feature a broad distribution of both the active and fully inactive I_2_ states capturing massive rearrangements of the key structural elements P-loop, A-loop, and αC-helix that are impossible to observe even in large-scale MD simulations. Consistent with the NMR experiments^40^ we demonstrate that introduction of the activating gate-keeper mutation T334I in the G269E/M309L/T334I and M309L/L320I/T334I triple ABL mutants can lead to the reverse shift by moderate the populations of the inactive states and promoting the dominant active ABL form. The results also highlight the inherent difficulties in accurately reproducing the mutation-induced relative populations of the ABL states, as AF2 predictions are intrinsically biased towards the ground ABL state. To facilitate physics-based analysis and interpretation of the AF2 predictions, we also perform a network-based allosteric analysis of the ABL structures and AF2-generated ensembles. Through this analysis we infer that functional ABL mutations can directly target or be structurally proximal to allosteric hotpots of long-range interactions. These results underscore that learning specific patterns of coevolutionary signals together with attention-based learning of allosteric couplings across homologous folds may augment the predictive abilities of AF2-based methods.

## Materials and Methods

### MSA Shallow Subsampling Adaptation of AF2

Structural prediction of the ABL kinase states were carried out using AF2 framework^1,2^ within the ColabFold implementation^46^ using a range of MSA depths and MSA subsampling.^18^ The MSAs were generated using the MMSeqs2 library^47,48^ using the ABL1 sequence from residues 240 to 440 as input. We used *max_msa* field to set two AF2 parameters in the following format: *max_seqs:extra_seqs*. These parameters determine the number of sequences subsampled from the MSA (*max_seqs* sets the number of sequences passed to the row/column attention track and *extra_seqs* the number of sequences additionally processed by the main evoformer stack). The default MSAs are subsampled randomly to obtain shallow MSAs containing as few as five sequences. This parameter is in the format of *max_seqs:extra_seqs* which decides the number of sequences subsampled from the MSA. *Max_seq* determines the number of sequences passed to the row/column attention matrix at the front end of the AF2 architecture, and *extra_seqs* sets the number of extra sequences processed by the Evoformer stack after the attention mechanism. The lower values encourage more diverse predictions but increase the number of misfolded models. We explored the following parameters: *max_seq, extra_seq*, number of seeds, and number of recycles. We ran simulations with *max_seqs:extra_seqs* 16:32, 32:64, 64:128. 128:256, 256:512 and 512:1024 values and report the results at *max_seqs:extra_seqs* 16:32 that produced the greatest diversity. We additionally manipulated the *num_recycles* parameters to produce more diverse outputs.

To generate more data, we set *num_recycles* to 12, which produces 14 structures starting from recycle 0 to recycle 12 and generating a final refined structure. Recycling is an iterative refinement process, with each recycled structure getting more precise. AF2 makes predictions using 5 models pretrained with different parameters, and consequently with different weights. Each of these models generates 14 structures, amounting to 70 structures in total. We then set the *num_seed* parameter to 1. This parameter quantifies the number of random seeds to iterate through, ranging from random_seed to random_seed+num_seed. We also enabled the use_dropout parameter, meaning that dropout layers in the model would be active during the time of predictions. This further increases variability within predictions.

### AF2 Adaptation Using Sequence Scanning with Shallow Subsampling

The initial input for the full sequence randomized alanine scanning is the original full native sequence. This technique utilizes an algorithm that iterates through each amino acid in the native sequence and randomly substitutes 5-15% of the residues with alanine, to simulate random alanine substitution mutations.^46^ The algorithm substitutes residue with alanine at each position with a probability randomly generated between 0.05 and 0.15 for each sequence position. We ran this algorithm multiple times (∼10-50) on the full sequences for each mutant, resulting in a multitude of distinct sequences, each with different frequency and position of alanine mutations. MSAs are then constructed for each of these mutated sequences using the alanine-scanned full-length sequences as input for the MMSeqs2 program.^47,48^ The AF2 shallow MSA methodology is subsequently employed on these MSAs to predict protein structures as described previously. A total of 70 predicted structures were generated from 12 recycles per model. In addition to randomized alanine sequence scanning of the complete sequence, we also examined several variations of this approach with targeted alanine masking of the ABL sequence space. In particular, we probed the effects of random alanine masking of sequence positions in the A-loop (resides 398-421) that is critical for conformational change between the active and inactive ABL forms. For each of these targeted alanine making experiments, we generate 10 alanine scanned sequences, each with different frequency and position of alanine mutations in the respective A-loop and C-terminal loop regions.

### Statistical and Structural Assessment of AF2-Generated Models

AF2 models were ranked by Local Distance Difference Test (pLDDT) scores (a per-residue estimate of the prediction confidence on a scale from 0 to 100), quantified by the fraction of predicted Cα distances that lie within their expected intervals. The values correspond to the model’s predicted scores based on the lDDT-Cα metric which is a local superposition-free metric that assesses the atomic displacements of the residues in the predicted model.^1,2^ Models ranked in the top five were compared to the experimental structure using structural alignment tool TM-align, an algorithm for sequence-independent protein structure comparison, to assess and compare the accuracy of protein structure predictions.^49^ TM-align involves optimizing the alignment of residues, refining the alignment through dynamic programming iterations, superposing the structures based on the alignment, and finally, calculating the TM-score as a quantitative measure of the overall accuracy of the predicted models. TM-align calculates the TM-score as the measure of overall accuracy of the prediction. An optimal superposition of the two structures is then built and TM-score is reported as the measure of overall accuracy of prediction for the models.^49^ TM-score ranges from 0 to 1, where a value of 1 indicates a perfect match between the predicted model and the reference structure. When TM-score > 0.5 implies that the structures share roughly the same fold. TM-score > 0.5 is often used as a threshold to determine if the predicted model has a fold similar to the reference structure. If the TM-score is above this threshold, it suggests that the predicted structure and the reference structure have a significant structural resemblance. Distributions of TM-Scores were calculated in terms of the averages of the highest TM-Scores for AF2 models compared to PDB structures of the same kinases for each MSA depth and structural conformation.

### Protein Structure Network Analysis

A graph-based representation of protein structures^50^ is used to represent residues as network nodes and the inter-residue edges to describe non-covalent residue interactions. The weights of the network edges in the residue interaction networks are determined by dynamic residue cross-correlations obtained from MD simulations^51^ and coevolutionary couplings between residues measured by the mutual information scores.^52^ The construction of the residue interaction network is described in detail in our previous studies.^53–55^ In brief, we summarize here the details as follows. The edge lengths in the network are obtained using the generalized correlation coefficients *R_MI_(x_i_, x_j_)* associated with the dynamic correlation and mutual information shared by each pair of residues. The length (i.e. weight) *w_ij_ = -log[R_MI_ (x_i_, x_j_)]* of the edge that connects nodes *i* and *j* is calculated from the corresponding generalized correlation coefficient between these nodes.^56^ The ensemble of shortest paths is determined from matrix of communication distances by the Floyd-Warshall algorithm.^57^ Network graph calculations were performed using the python package NetworkX.^58^ Using the constructed protein structure networks, we computed the residue-based betweenness parameter. The betweenness of residue *i* is defined to be the sum of the fraction of shortest paths between all pairs of residues that pass through residue *i*:

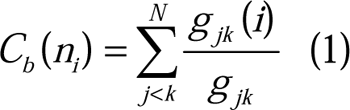

where *g_jk_* denotes the number of shortest geodesics paths connecting *j* and *k*, and *g _jk_* (*i*) is the number of shortest paths between residues *j* and *k* passing through the node *n_i_*.

The Girvan-Newman algorithm^59–61^ is used to identify local communities and optimize modularity of the interaction network. In this approach, edge centrality (edge betweenness) is defined as the ratio of all the shortest paths passing through a particular edge to the total number of shortest paths in the network. To characterize global bridges from a community structure, we introduce a community bridgeness metric similar to Rao-Stirling index.^62–64^ This parameter uses as input a prior categorization of the nodes into distinct communities:

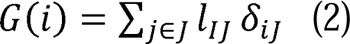

where the sum is over communities *J* (different from the community of node *i*, denoted as *I*), δ_iJ_ is equal 1 if there is a link between node *i* and community *J* and 0 otherwise. *l_ij_* corresponds to the effective distance between community *I* and community *J* as measured by the inverse of the number of links between them. All topological measures were computed using the python module the python package NetworkX^58^ and also Cytoscape platform for network analysis.^65^ The ModuLand program within the Cytoscape platform^66,67^ was also adapted to determine a hierarchical network structure and compute residue bridgeness of the identified communities in the dynamic residue interaction networks. The RING program^68,69^ was used to generate the initial residue interaction networks.

## Results and Discussion

### Atomistic Modeling of the Conformational Ensembles of the ABL Kinase Mutants Using Shallow MSA Subsampling AF2 Approach: Emerging Bias Towards the Active ABL form

Conformational transitions between kinase states are orchestrated by three conserved structural motifs in the catalytic domain: the αC-helix, the 400-DFG-402 motif in ABL (DFG-in, active; DFG-out, inactive), and the activation loop (A-loop open, active; A-loop closed, inactive). The conserved His-Arg-Asp (HRD) motif in the catalytic loop and the DFG motif are coupled with the αC-helix to form conserved intramolecular networks termed regulatory spine (R-spine) and catalytic spine (C-spine). The R-spine in ABL includes M309 from the C the αC-helix, L320 from the β4-strand, F401 of the DFG motif in the beginning of the A-loop, H380 of the HRD motif in the catalytic loop, and D440 of the αF-helix (Figure 1A-C). The R-spine subnetwork is fully assembled in the active ABL kinase (Figure 1A), but it becomes decoupled in the inactive state I_1_ (Figure 1B) and is completely broken in the inactive state I_2_ (Figure 1C). The C-spine is comprised of hydrophobic residues (V275, A288, L342, C388, L389, V336, S457, and I451) connects the kinase lobes anchoring catalytically important sites to the C-terminus of the αF-helix. The NMR ensemble of the active conformations (pdb id 6XR6) is characterized by the “αC-in” position and stable DFG-in orientation (Figure 1D), while in the inactive I_1_ state (pdb id 6XR7) the αC helix moves to the intermediate αC-out position and the DFG motif is flipped 180°, with respect to the active conformation (Figure 1E). In the inactive I_2_ state, the regulatory DFG motif assumes a distinct “out” conformation and the A-loop swings to a fully closed conformation (Figure 1C). The original structural studies remarked that while Abl I_2_ state adopts a fully inactive conformation, the inactive intermediate I_1_ state may lie in the pathway between the active and the I_2_ state.^40^

**Figure 1.**
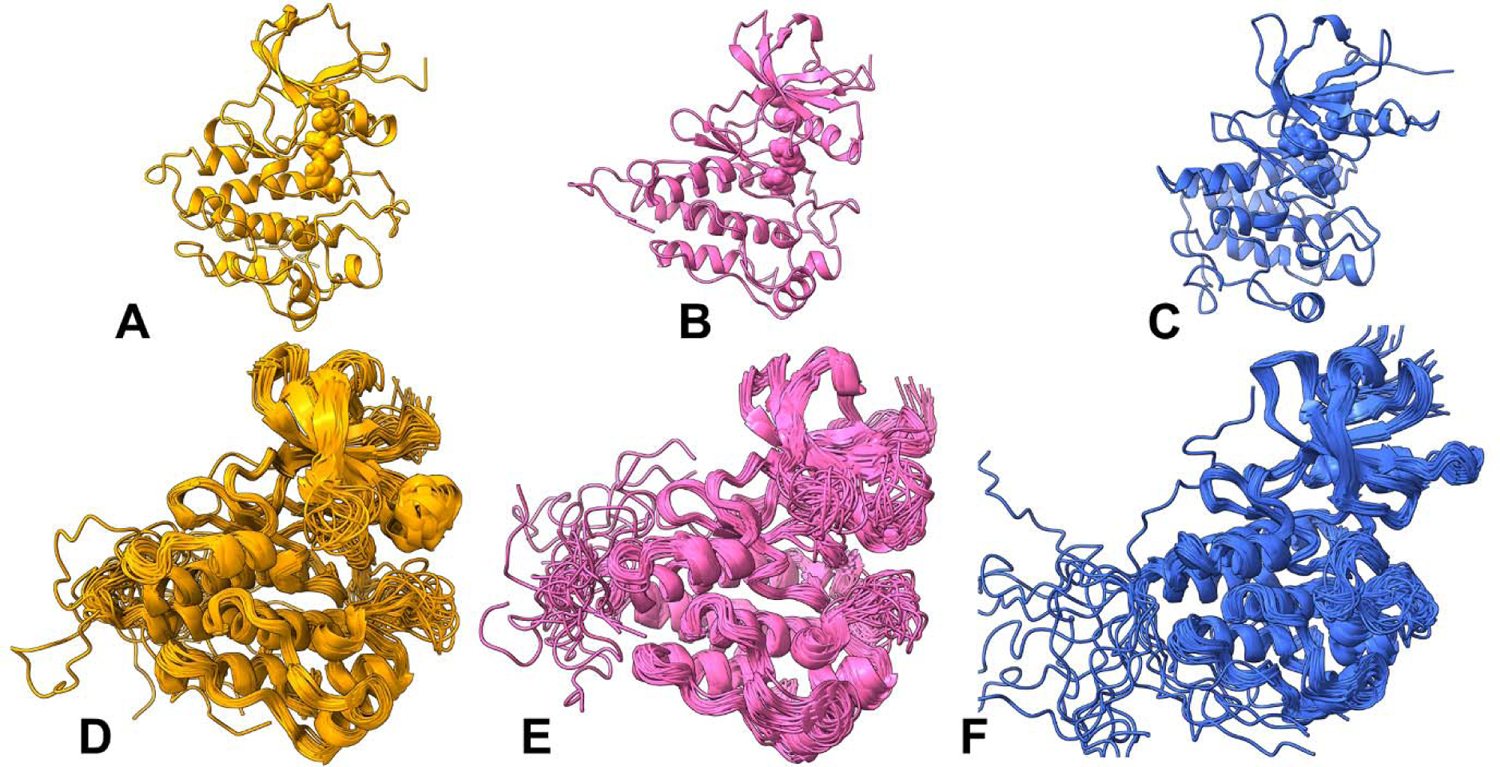
The thermodynamically stable fully active ground state of the ABL kinase domain (orange ribbons, pdb id 6XR6) (A), the inactive state I_1_ (pink ribbons pdb id 6XR7) (B) and the closed inactive state I_2_ (blue ribbons, pdb id 6XRG) (C). The R-spine residues M309, L320, H380, F401 in the active state and inactive state I_1_ are shown in spheres respectively. The R-spine positions in the inactive state I_2_ are L309, L320, H380, F401 and are shown in blue spheres. The NMR solution structural ensembles of the active ground state of the ABL kinase domain (pdb id 6XR6) (D), the inactive state I_1_ (pdb id 6XR7) (E) and the closed inactive state I_2_ (pdb id 6XRG) (F). The NMR ensembles are depicted in orange, pink and blue ribbons respectively for the active and inactive I_1_ and I_2_ ABL forms.

NMR studies established that M309L mutation in the R-spine increases the population of I_1_ and I_2_ to 10% and 35% respectively, while double M309L/L320I substitution switches the ensemble towards the I_2_ state (∼82%) thus deactivating the kinase.^40^ The ABL mutant M309L/H415P shifts the equilibrium towards I_1_ without eliciting other structural changes, while T408Y/H415P, increases the population of I_2_ state.^40^ G269E modification in the P-loop is Imatinib-resistant mutation acts by enforcing the kinked P-loop conformation observed in the active state and destabilizing the stretched conformation in the I_2_ state. At the same time, triple ABL mutant G269E/M309L/T408Y that involves modifications in the P-loop (G269E), R-spine (M309L) and A-loop (T408Y) shifts the equilibrium from the active state towards the I_2_ state at ∼85% population, while activating mutation T334I can moderate the populations of the inactive states in the G269E/M309L/T334I and M309L/L320I/T334I triple mutants.^40^

Hence, according to the NMR data, regulatory mutations can promote unique structural and functional effects by shifting the equilibrium between the thermodynamically dominant active ABL form and the inactive I_1_ and I_2_ states (Supporting Information, Figure S1). We employed several different AF2 adaptations, including shallow MSA subsampling approach, to predict structures and conformational ensembles for a series of state-switching double ABL mutants M309L/L320I, M309L/H415P, T408Y/H415P and triple mutants G269E/M309L/T408Y, G269E/M309L/T334I, M309L/L320I/T334I (Supporting Information, Figure S1) that have unique effects on the conformational equilibrium and can induce large conformational transformations between the active and inactive ABL ensembles. In addition, we compared the predictive performance of AF2 methods with the recently proposed in our laboratory AF2 adaptation in which randomized alanine sequence scanning of the entire protein sequence or masking of specific functional regions is combined with the MSA subsampling.^45^ By using different AF2 adaptations we examined the predictive ability and limitations of these tools to detect functional states and characterize mutation-specific shifts in conformational ensembles of the ABL mutants. First, we analyzed the AF2 predictions for ABL mutants using shallow MSA approach. The AF2-produeced MSAs are summarized as a heatmap indicating all sequences mapped to the input sequences (Supporting Information, Figure S2). The relative coverage of the sequence with respect to the total number of aligned sequences is shown indicating the reduced sequence identity to query for the highly flexible N-lobe regions, while a statistically significant sequence coverage was seen for the kinase domain core regions and C-lobe residues (Supporting Information, Figure S2). We found that shallow MSA subsampling predictions converged predominantly to the active ABL conformation for all examined ABL mutants, showing high confidence pLDDT values (Supporting Information, Figure S3). Most of the ABL kinase core regions featured pLDDT ∼80-100, while the A-loop (residues 395-421) displayed moderately reduced pLDDT values ∼65-85 and low pLDDT values only for highly flexible N-lobe regions (Supporting Information, Figure S3). The pLDDT profiles for the top models are very similar for all studied mutants but showed a moderately increased variability among models for the triple mutants (Supporting Information, Figure S3D-F). Some of the low confidence pLDDT values corresponded to disordered N-terminal residues that were revealed in the NMR ensembles. To gain a quantitative insight into the AF2 predictions, we constructed the pLDDT density distribution for the predicted conformational ensembles of the ABL mutants (Figure 2). The dominant peaks at pLDDT ∼85-90 and several minor peaks for pLDDT∼70-75 are present for all mutants, indicative of similar conformational ensembles produced by shallow MSA approach for all mutants. Instructively, shallow MSA subsampling predictions reflected conformational heterogeneity of the active state that is exemplified in appreciable fluctuations of the open A-loop conformations.

**Figure 2.**
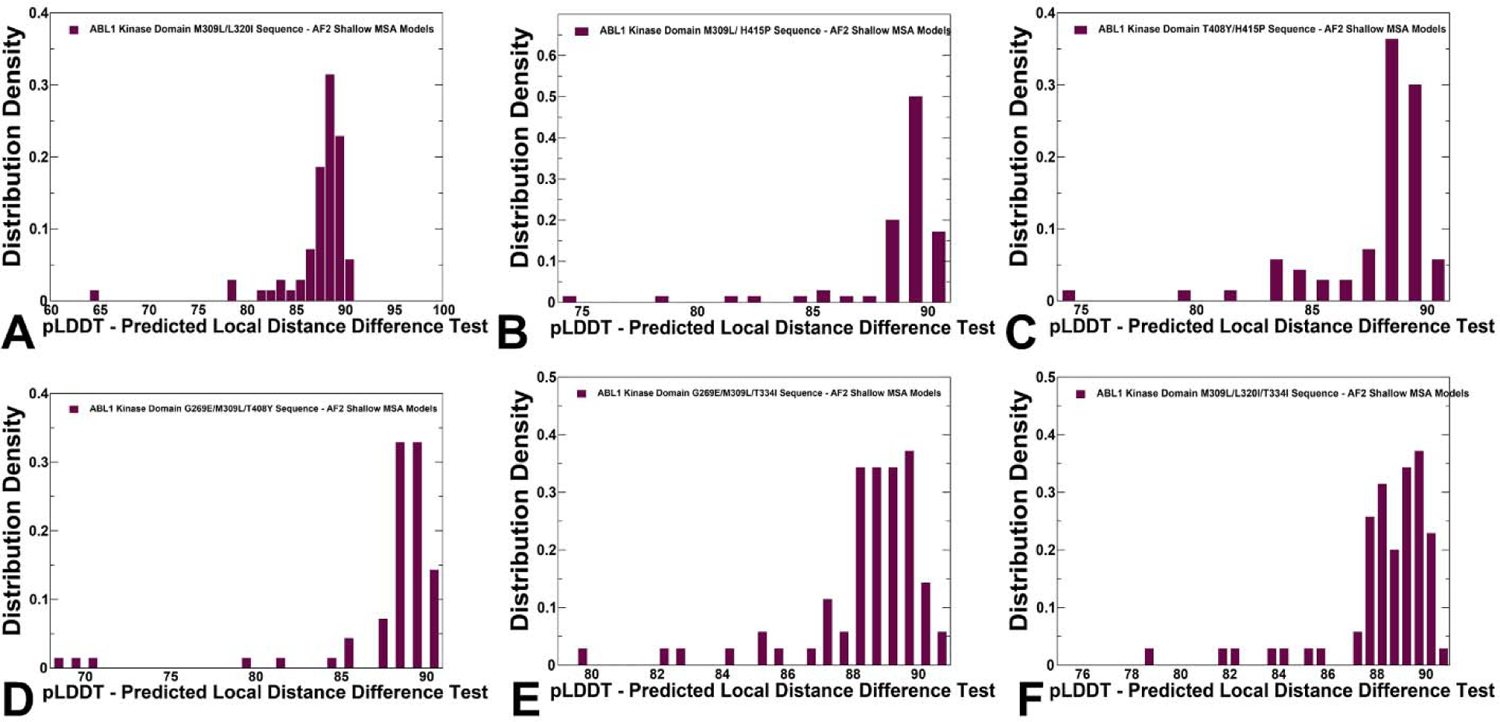
The statistical analysis of the shallow MSA depth ensemble models for the ABL kinase mutants. The density distributions of the pLDDT estimates for the predicted models in conformational ensembles of the M309L/L320I mutant (A), M309L/H415P mutant (B), H415P/T408Y mutant (C), G269E/M309L/T408Y mutant (D), G269E/M309L/H415P mutant (E) and M309L/L320I/T334I mutant (F). The pLDDT structural model estimate of the prediction confidence is on scale from 0 to 100. The pLDDT values within 70-90 range, suggest high confidence of the predictions, pLDDT values ∼ 50-70 signal lower confidence and must be treated with caution, while pLDDT < 50 may be a reasonably strong predictor of disorder.

The TM scores of the AF2-predicted conformations showed narrow peaks of TM values ∼0.85-0.88 with respect to the active ABL conformation, indicating that the overwhelming majority of the predicted conformations are structurally similar to the active ABL form (Supporting Information, Figure S4). We also computed the RMSD distributions of the predicted ensembles with respect to the active and inactive experimental structures (Figure 3). The RMSD densities showed a clear separation between the RMSDs computed with respect to different ABL states, particularly highlighting the dominant peaks for RMSDs < 1.0 Å relative to the active form (Figure 3A-C). We observed contributions of conformations that are similar to the intermediate inactive I_1_ form (RMSD ∼ 2.0Å) but these similarities largely reflect the intermediate nature of the inactive I_1_ state in which the A-loop remains in an open conformation and the N-terminal part of the A-loop is very similar in the active and I_1_ states. However, NMR ensemble for the I_1_ form features the DFG motif being flipped 180° with respect to the active conformation and adopting DFG-out position.

**Figure 3.**
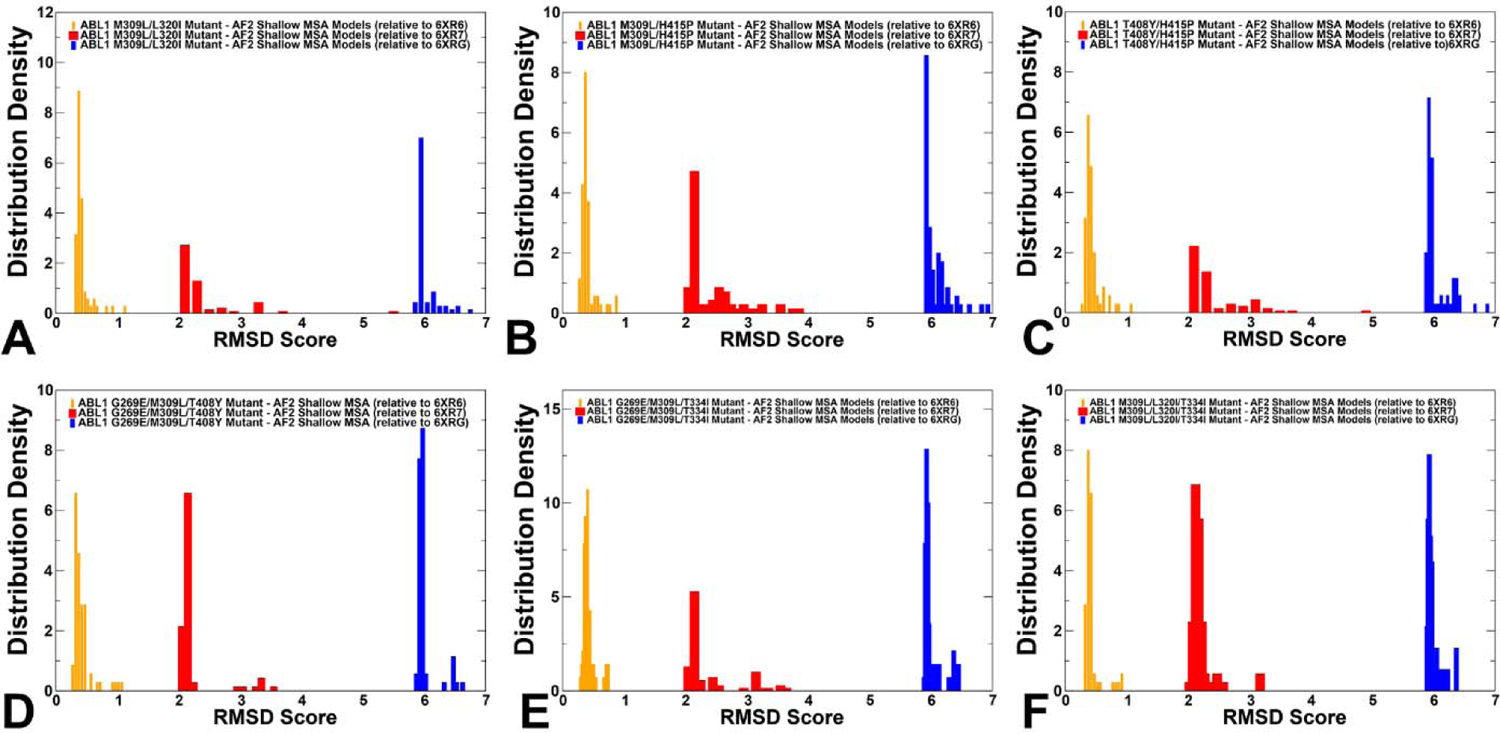
The analysis of AF2 predictions of the conformational ensembles for ABL mutants using shallow MSA subsampling approach. The density distributions of the RMSD values for predicted conformational ensembles of the M309L/L320I mutant (A), M309L/H415P mutant (B), H415P/T408Y mutant (C), G269E/M309L/T408Y mutant (D), G269E/M309L/H415P mutant (E) and M309L/L320I/T334I mutant (F). The distribution density of RMSD scores for the AF2-predicted conformations are shown relative to the active ABL state (orange filled bars) and the inactive states I_1_ (red filled bars) and I_2_ (blue filled bars).

Structural mapping of the conformational ensembles (Figure 4) showed moderate perturbations of the A-loop in most of the mutants with the exception of M309L/L330I and G269E/M309l/T408Y where more significant fluctuations were seen without changing the extended open arrangement of the A-loop. The alignment of the DFG conformations showed that both DFG-in and DFG-out conformations which is characteristic of I_1_ can be found only for M309L/L320I double mutant and G269E/M309L/T408Y mutants (Figure 4A,D).

**Figure 4.**
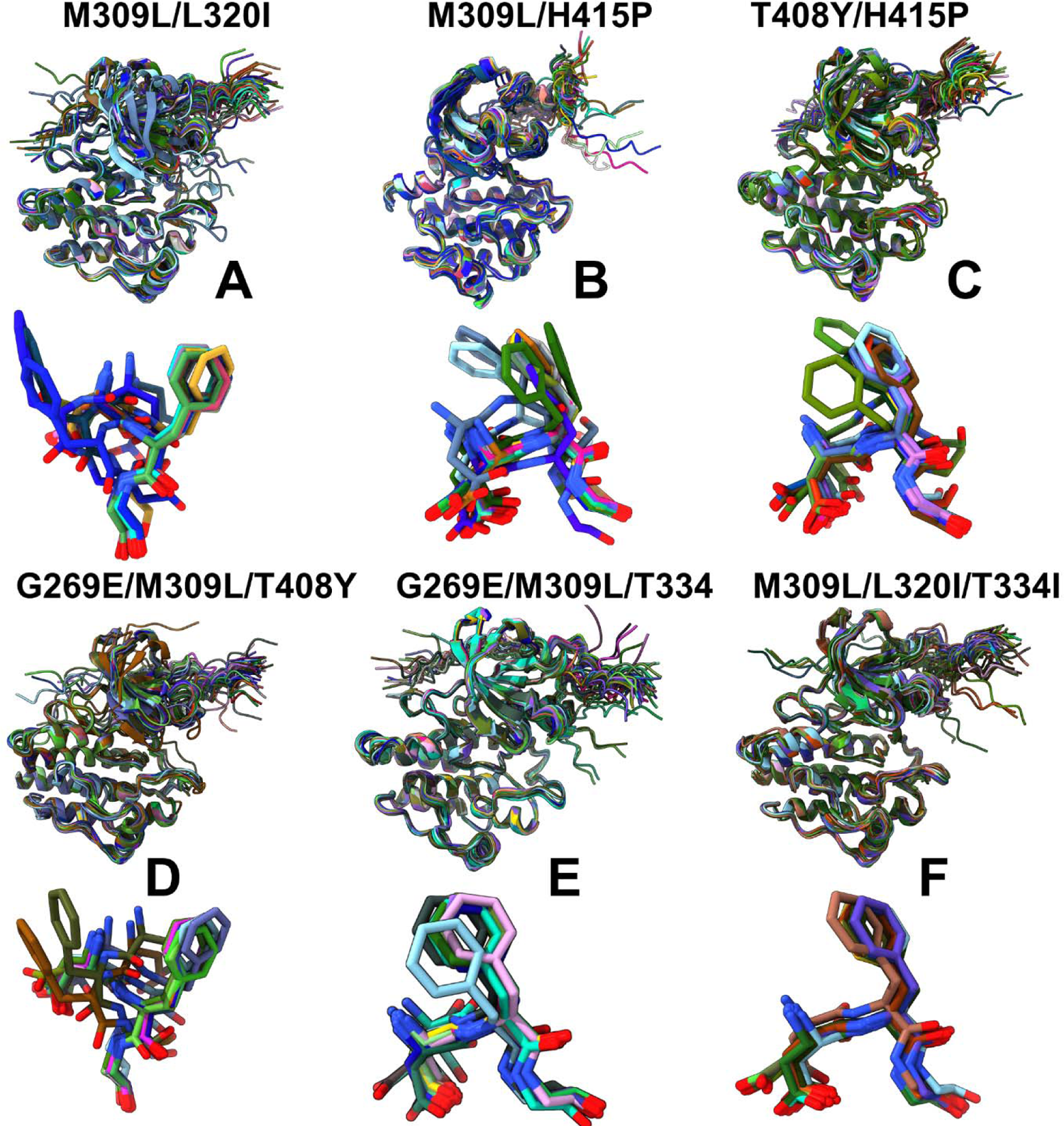
Structural alignment of the AF2-predicted ensembles using shallow MSA subsampling approach. Structural overlay of the kinase catalytic domain conformations and DFG motif from the AF2-predicted conformational ensemble are shown for the M309L/L320I mutant (A), M309L/H415P mutant (B), H415P/T408Y mutant (C), G269E/M309L/T408Y mutant (D), G269E/M309L/H415P mutant (E) and M309L/L320I/T334I mutant (F).

In these mutants, DFG motif samples intermediate “out” positions with F401 pointing upward. DFG conformations in triple mutants sharing T334I mutation conform to the active DFG-in position with relatively modest fluctuations of the open A-loop (Figure 4E,F). To characterize conformational changes in the A-loop, we also computed the RMSD densities for the A-loop resides only (Figure 5). This analysis confirmed that the majority of the ensemble conformations conforms to the active ABL form, revealing larger fluctuations of the active A-loop and emergence of the conformations with A-loop similar to the inactive I_1_ state (RMSD ∼ 2.8-3.5 Å) for the M309L/L320I and M309L/H415P double mutants (Figure 5A,B).

**Figure 5.**
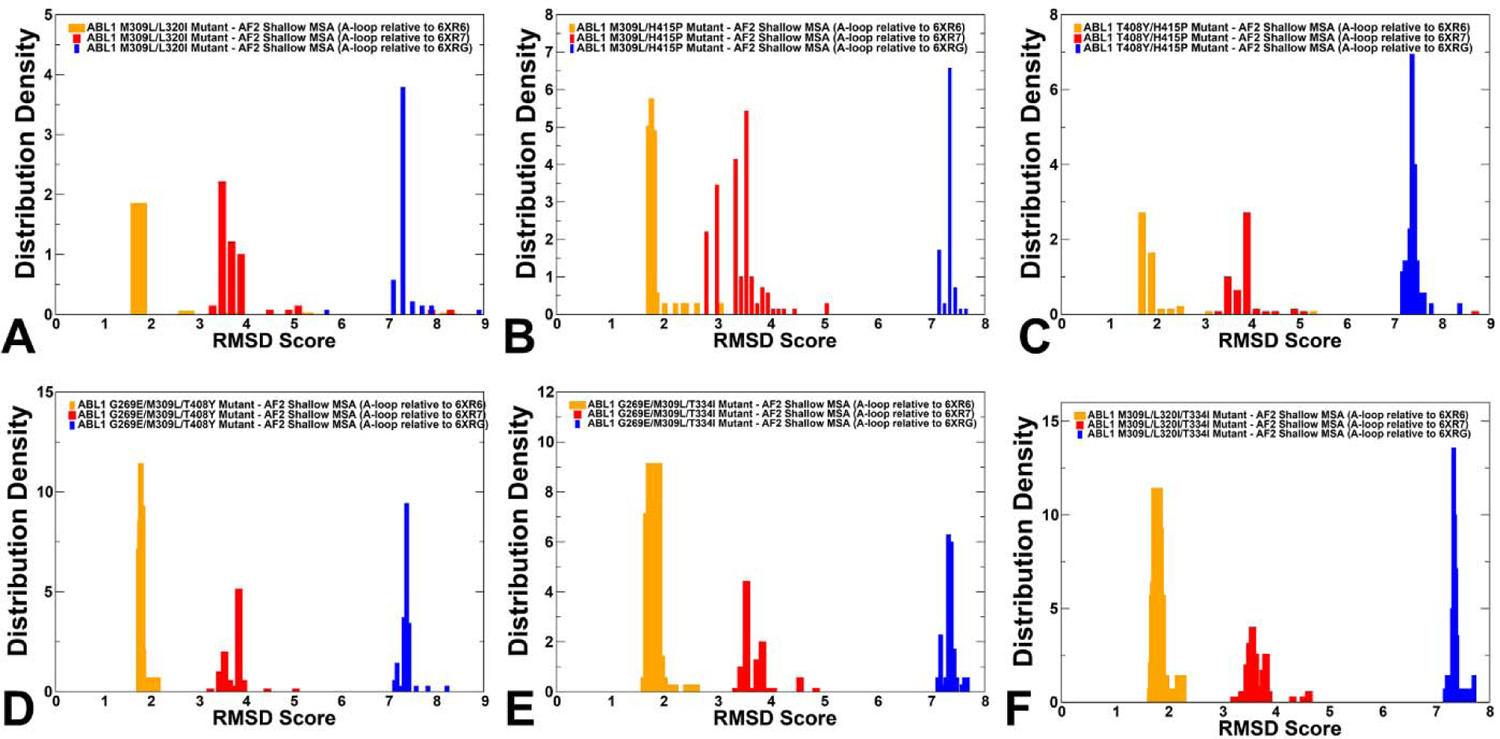
The analysis of AF2 predictions of the conformational ensembles for ABL mutants using shallow MSA subsampling. The density distributions of the RMSD values for the A-loop (residues 395-421) in the predicted ensembles are shown for M309L/L320I (A), M309L/H415P (B), H415P/T408Y (C), G269E/M309L/T408Y (D), G269E/M309L/H415P (E) and M309L/L320I/T334I mutants (F). The distribution density of RMSD scores of the A-loop for the AF2-predicted conformations are shown relative to the active ABL state (orange filled bars) and the inactive states I_1_ (red filled bars) and I_2_ (blue filled bars).

NMR data showed that the H415P mutation can destabilize the I_2_ state and increase stabilization of the I_1_ state.^40^ The shallow MSA AF2 predictions may reflect the increased heterogeneity of the A-loop induced by mutations in the A-loop (T408Y and H415P), in some conformations but the overall predicted ensembles are largely dominated by the active ABL states. The RMSD distributions for the triple mutants, particularly for G269E/M309L/T334I and M309L/L320I/T334 showed the dominance of the active ABL conformations with narrow RMSD peaks ∼ 1.5-2.0 Å from the active structure (Figure 5D-F). In this context, it is important to note that experimentally the presence of the activating T334I gate-keeper mutation promotes formation of the active form and counteracts the effect of other mutations.

Principal Component Analysis (PCA) of the AF2-predicted ensembles and experimental NMR ensembles^40^ of the active ABL state, inactive I_1_ and inactive I_2_ forms revealed important similarities and differences, pointing out to sampling capabilities of the shallow MSA subsampling approach (Figure 6A-C). The generated models from all methods were processed to exclude models with a pLDDT < 60-70. MDAnalysis library for PCA of the structural ensembles created the trajectories of all the generated structures and then PCA algorithm was employed to project the respective conformational ensembles on two principal components (Figure 6). The AF2-predicted ensembles for the ABL mutants are confined to some regions of the conformational space relevant to the functional states and cannot provide accurate mapping of the NMR ensembles for the inactive ABL states. However, PCA of the conformational ensembles for M309L/L320I and M309L/H415I double mutants displayed a broader coverage of the configurational space, particularly indicating sampling of the inactive-like states. According to the NMR data^40^, these mutants shift the equilibrium away from the active form, and M309L/H415P shifts the equilibrium towards I_1_ without eliciting other structural changes. Hence, the analysis of the AF2 predictions using shallow MSA depth adaptation revealed only a limited variability of the regulatory DFG-in conformation that remained confined to its active form. Although subsampling of MSA may increase conformational heterogeneity around the ground active ABL state, this approach cannot readily generate inactive ABL conformations that become more dominant in the ABL double mutant M309L/L320I and triple mutant G269E/M309L/T408Y.

**Figure 6.**
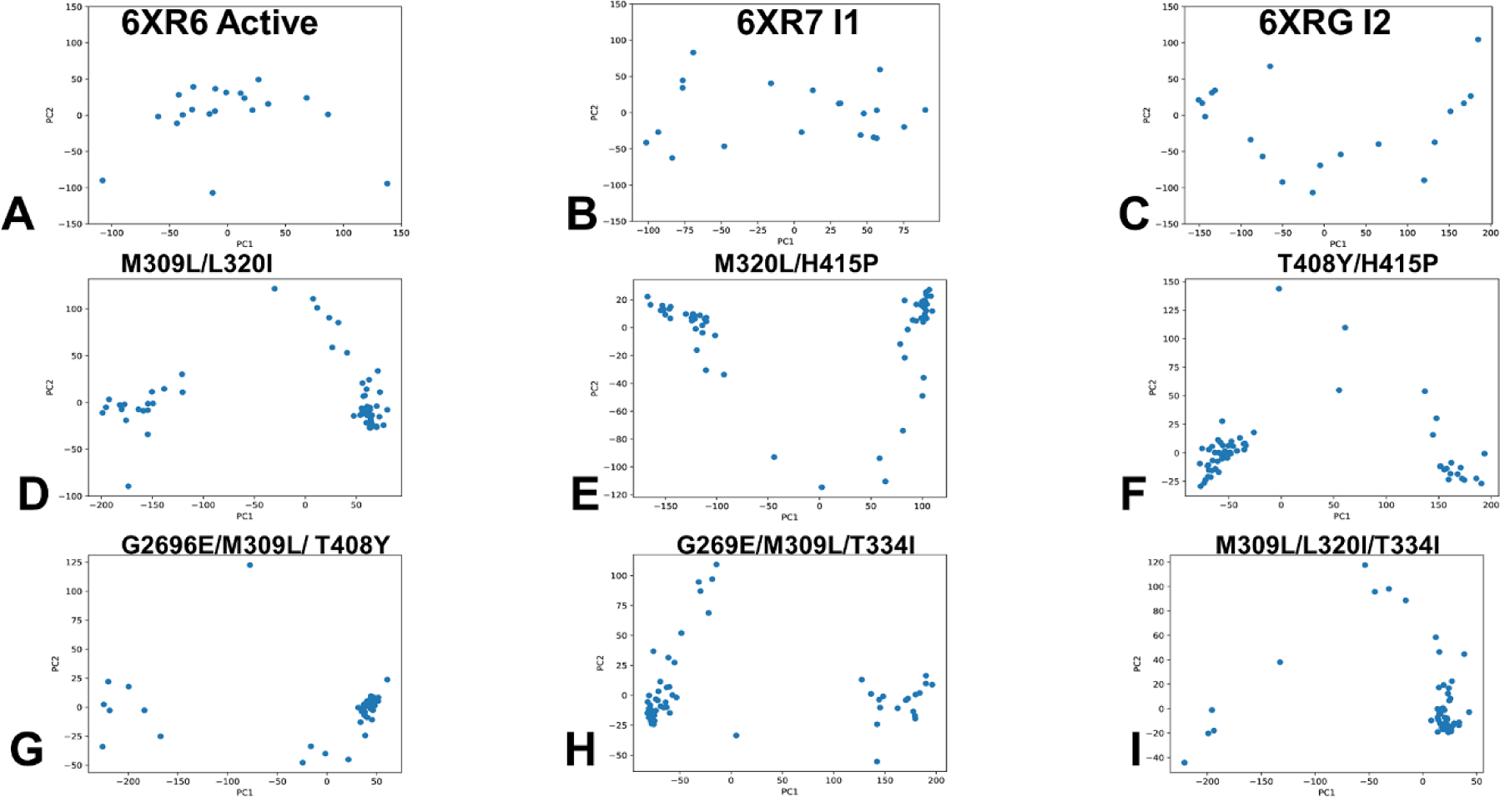
PCA of the NMR ensembles for the active ABL structure (A), inactive I_1_ structure (B) and inactive I_2_ structure (C). PCA of the generated ensembles for ABL mutants using shallow MSA subsampling are shown for the M309L/L320I mutant (D), M309L/H415P mutant (E), H415P/T408Y mutant (F), G269E/M309L/T408Y mutant (G), G269E/M309L/H415P mutant (H) and M309L/L320I/T334I mutant (I).

### Alanine Sequence Scanning Combined with Shallow MSA Subsampling Can Detect Allosteric ABL States and Capture Population Shifts in State-Switching ABL Mutants

We employed recently developed randomized alanine scanning adaptation of the AF2 methodology in which the algorithm operates first on the pool of sequences and iterates through each amino acid in the native sequence to randomly substitute 5-15% of the residues with alanine, thus emulating random alanine mutagenesis.^45^ The algorithm substitutes residue with alanine at each position with a probability randomly generated between 0.05 and 0.15 for each sequence position. In the proposed protocol, randomized alanine sequence scanning is followed by construction of corresponding MSAs and then by AF2 shallow subsampling applied on each of these MSAs. To probe differences between randomized alanine scanning of the entire protein sequence and targeted random scanning of specific kinase regions, we also examined several variations of this approach with targeted alanine masking of the ABL sequence space. Random alanine masking of sequence positions in regions involved in conformational changes included the A-loop (resides 398-421), the αC-helix (residues 300-311) and the C-lobe regions. For each of these targeted alanine making experiments, we generate 50 alanine scanned sequences, each with different frequency and position of alanine mutations in the respective regions. By combining manipulation of sequence variations with the MSA subsampling, we probed the effects of randomized alanine scanning in different kinase domain regions on accuracy and diversity of the predicted AF2 ensembles.

We analyzed the RMSD values computed for the predicted ABL structure with respect to the active and inactive experimental ABL structures (Figure 7). The results showed a significant overlap in the distributions for M309L/L320I (Figure 7A), M309L/H415P (Figure 7B) and G269E/M309L/T408Y triple mutant (Figure 7C). This implies that the predicted conformational ensembles for these ABL mutants also featured appreciable populations of both I_1_ and I_2_ forms along with the active form. Strikingly, we found that for the M309L/L320I and G269E/M309L/T408Y mutants the population of the fully inactive I_2_ form may become highly significant (Figure 7A,D). This is consistent with the experimentally observed shifts towards the inactive state for these ABL mutants.^40^ Notably, the RMSD distribution of predicted conformations with respect to the inactive I_2_ conformation showed a peak at RMSD ∼ 2.0 Å for these mutants, while also reproducing population of the active ABL conformations with RMSDs ∼ 1.0 Å from the active structure (Figure 7A-D).

**Figure 7.**
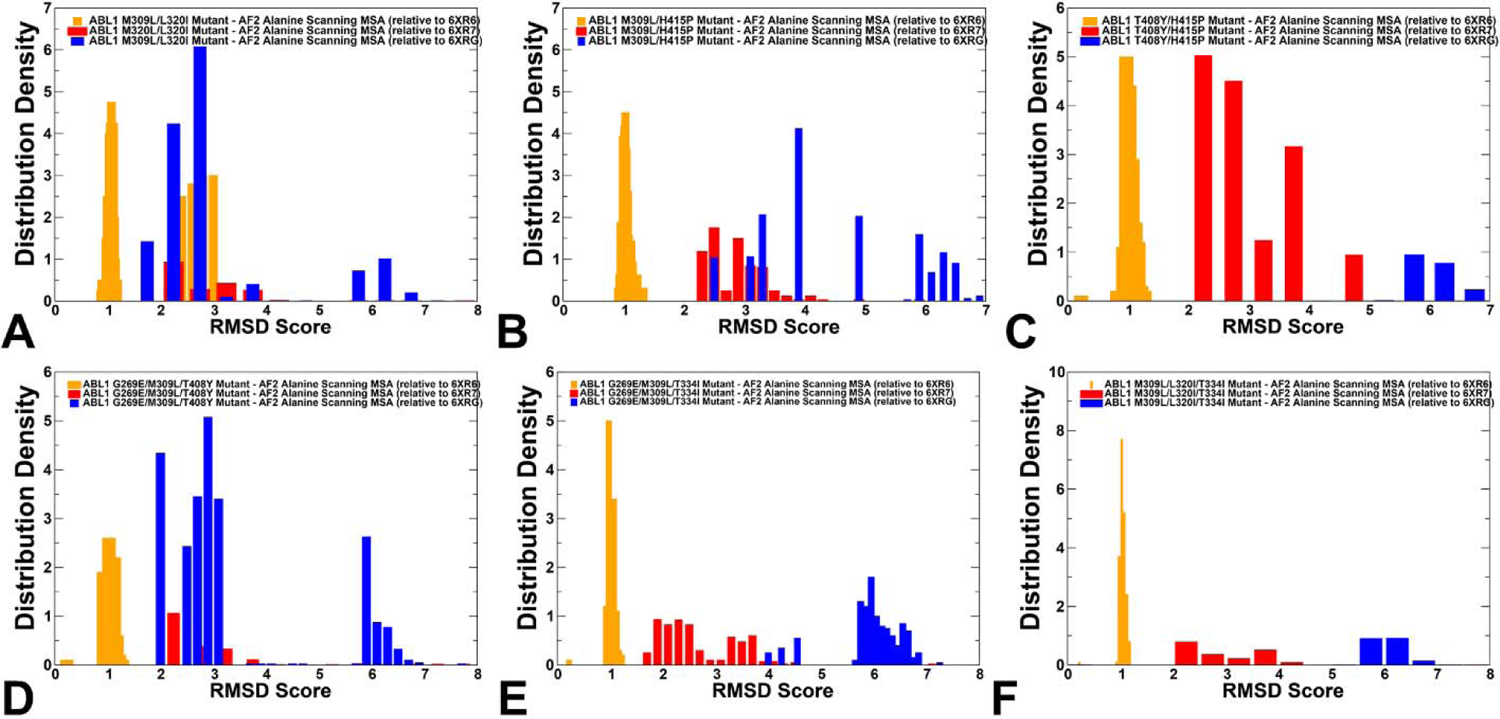
The analysis of AF2 predictions using randomized alanine sequence scanning approach. The distribution density of RMSD scores for the AF2-predicted ABL conformational ensembles of the M309L/L320I mutant (A), M309L/H415P mutant (B), H415P/T408Y mutant (C), G269E/M309L/T408Y mutant (D), G269E/M309L/H415P mutant (E) and M309L/L320I/T334I mutant (F). The distribution density of RMSD scores for the AF2-predicted conformational ensembles are shown relative to the active ABL state (orange filled bars) and the inactive states I_1_ (red filled bars) and I_2_ (blue filled bars).

These findings can be contrasted with the predictions using shallow MSA approach where the vast majority of the generated conformations for all ABL mutants converged towards the active ABL form and could not adequately capture the mutation-induced switching to the inactive I_2_ form. Although these findings cannot be directly compared with the NMR analysis of relative populations showing that the double M309L/L320I substitution switches the ensemble towards the I_2_ state (∼82%) and mutant G269E/M309L/T408Y shifts the equilibrium towards the I_2_ state at ∼85% population^40^, a combination of randomized alanine masking with shallow MSA showed markedly improved predictions of the conformational ensembles. The predictions produced a significant functionally relevant ensemble of the heterogeneous inactive I_1_ conformations in the M309L/H415P mutant. Noticeably, and consistent with the NMR experiments, the introduction of the activating T334I mutation in the triple mutants reduced presence of the inactive conformations, displaying a strong single peak for RMSDs ∼ 1.0 Å from the active structure (Figure 7E,F).

Structural mapping of the predicted ABL conformations illustrated a considerably increase conformational heterogeneity of structural ensembles, also highlighting important differences between ABL mutants (Figure 8). Indeed, we observed significant variations of the A-loop that sampled both the open (active) and closed (inactive) conformations in the M309L/L320I (Figure 8A), M309L/H415P (Figure 8B) and G269E/M309L/T408Y mutants (Figure 8D). An appreciable variability of the open A-loop conformations was seen for the double mutant T408Y/H415P signaling that predicted ensemble sampled the active (αC-in, DFG-in, A-loop open) and intermediate inactive I_1_ ABL states (αC-in, DFG-in/out, highly flexible open A-loop (Figure 8C). Interestingly, we noticed that conformational variability of the predicted ensemble for triple mutants G269E/M309L/T334I (Figure 8E) and M309L/L320I/T334I (Figure 8F) is largely confined to functional fluctuations around the active ABL form. While the N-lobe regions in these triple mutants displayed significant plasticity, the key functional regions remained in their active positions (αC-in, DFG-in, A-loop open). Hence, structural analysis of the predicted ensembles for G269E/M309L/T334I and M309L/L320I/T334I mutants further suggested that presence of the activating T334I mutation shared by these mutants may promote accumulation of the active conformations that dominate the population. These findings are consistent with the NMR studies, showing that introduction of T334I can reverse the population of G269E/M309L/T408Y mutant from predominantly inactive I_2_ state (with only 10% of active population) to a dominant active form (with 93% of the total population).^40^

**Figure 8.**
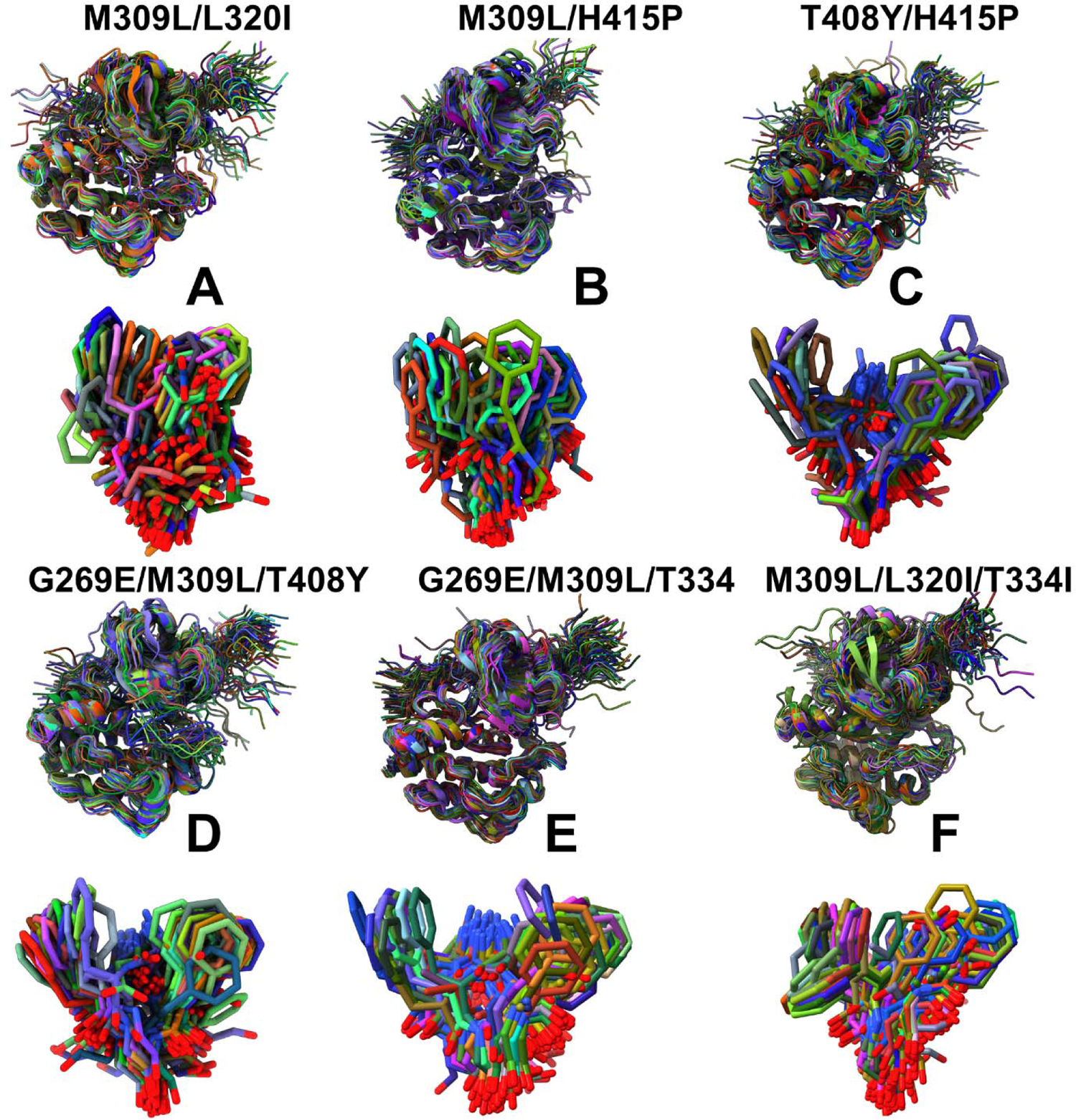
Structural alignment of the conformational ensembles for the kinase domain and DFG motif in the ABL mutants obtained using combination of alanine sequence scanning and shallow MSA subsampling. Structural overlay of the AF2-predicted conformational ensemble and DFG motifs are shown for the M309L/L320I mutant (A), M309L/H415P mutant (B), H415P/T408Y mutant (C), G269E/M309L/T408Y mutant (D), G269E/M309L/H415P mutant (E) and M309L/L320I/T334I mutant (F).

By projecting the AF2-produced DFG conformations, the functionally relevant positional variability of the DFG motif between active DFG-in position and inactive DFG-out conformations can be seen (Figure 8). The variability of the DFG motif is particularly exemplified by the observed movements of F401 residue that samples a large number of intermediate states between DFG-in and DFG-out flipped by 180° for all ABL mutants, In particular, we observed extensive sampling of the DFG-in and DFG-out positions for M309L/L320I mutant (Figure 8A), M309L/H415P (Figure 8B) and G269E/M309L/T408Y mutant (Figure 8D) that are known to shift populations of the ABL from the active to inactive I_2_ form. Consistent with our analysis, the population of the active DFG-in conformations remained significant in the triple mutants sharing activating gate-keeper mutation T334I (Figure 8E,F). The generated AF2 ensembles produced a significant population of the inactive I_1_ conformations that sample changes between αC-in and αC-out inactive positions as well as intermediate DFG-in to DFG-out orientations.

We also illustrated the predicted inactive I_2_ conformations for the ABL state-switching mutant M309L/L320I (Supporting Information, Figure S5). It should be noted that the population of these inactive states remains relatively diverse but share flipped A-loop conformation that switches from the open to closed conformation. The predicted conformations also featured some variability of the DFG-out position (Supporting Information, Figure S5), further highlighting the intrinsic difficulties of AF2 approached to precisely reproduce the hidden inactive forms of state-switching proteins. In this context, it is worth noting that the I_2_ state is a low-populated conformational state that resembles the Imatinib complex in the A-loop and DFG motif but is different in the αC helix and P-loop. The precise structural arrangement of these functional regions can be induced by the inhibitor binding, while our predictions are performed for the native and mutant ABL sequences devoid of any inhibitor information.

Of particular interest is a comparison of PCA plots for conformational ensembles produced by randomized alanine scanning (Figure 9) against PCA distributions obtained using shallow MSA approach (Figure 6). This analysis confirmed that the conformational ensembles generated using the proposed AF2 adaptation are considerably more diverse and enable more adequate sampling of functionally relevant states. Indeed, it can be seen that the distributions of sampled conformations for M309L/L320I (Figure 9A), M309L/H415P (Figure 9B) and triple mutant G269E/M309L/T408Y (Figure 9D) overlap with the NMR ensembles for both active and inactive states. It is particularly instructive to underscore the overlaps between the PCA distributions for these mutants and PCA for the NMR ensemble of the fully closed inactive I_2_ state. This analysis further confirms that the AF2-generated ensembles for these state-switching ABL mutants sample an appreciable population of the inactive form that is drastically different from the active form. The PCA analysis of the T408Y/H415P mutant (Figure 9C) displayed distribution that is highly reminiscent of the NMR ensemble highlighting preferences towards intermediate inactive I_1_ state. Finally, PCA distributions obtained for G269E/M309L/T334 and M309L/L320I/T334 triple mutants (Figure 9E.F) showed dense concentrations of conformations in the active-like conformations as can be judged from comparison with the PCA of the NMR ensemble of active conformations. This further confirms our findings suggesting that T334I mutation may promote shifts to a predominantly active form.

**Figure 9.**
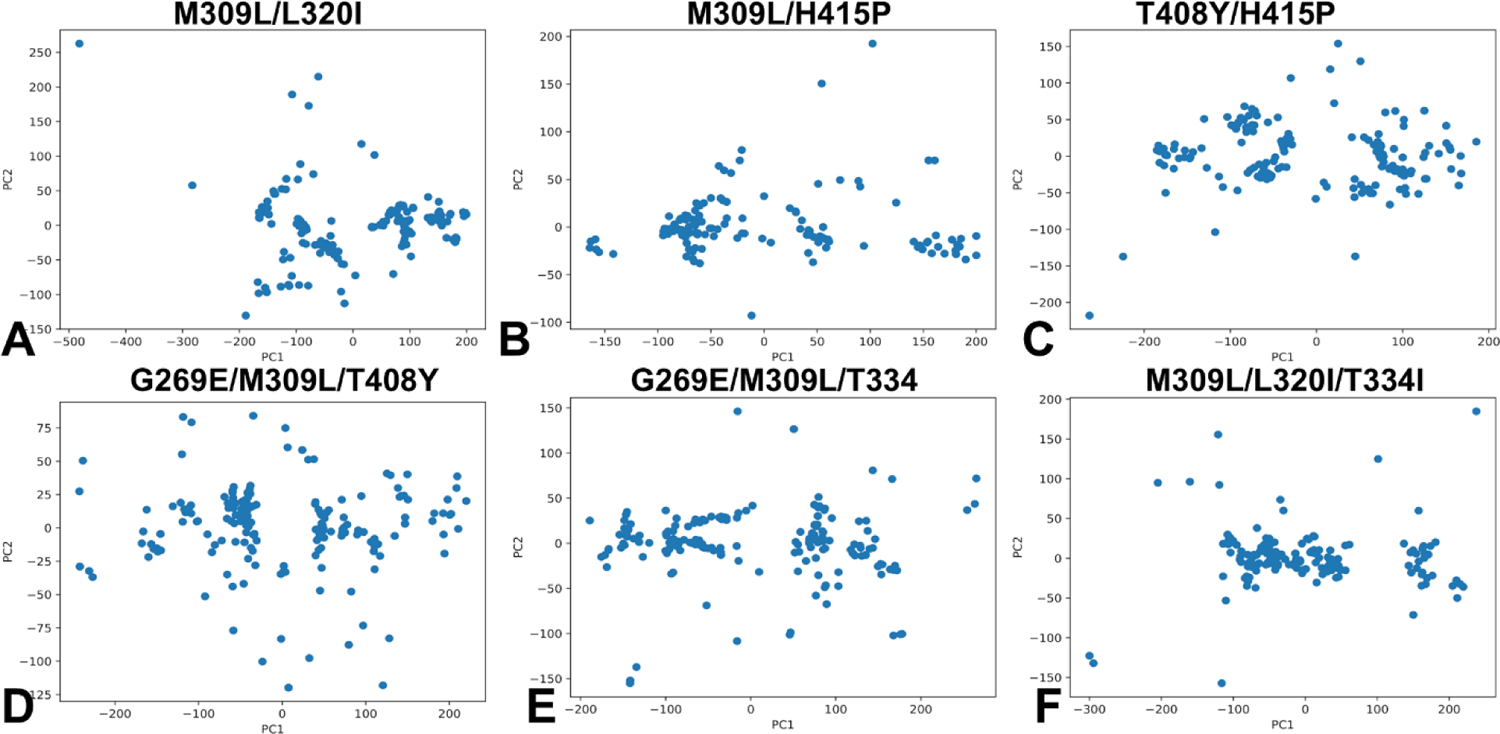
PCA of the generated ensembles for the ABL mutants using combination of alanine sequence scanning and shallow MSA subsampling. PCA plots are shown for the M309L/L320I mutant (D), M309L/H415P mutant (E), H415P/T408Y mutant (F), G269E/M309L/T408Y mutant (G), G269E/M309L/H415P mutant (H) and M309L/L320I/T334I mutant (I).

Together with our previous analyses, the results demonstrate the ability of the proposed approach to capture physically significant population shifts and produce distinct functional conformational clusters that are associated with the active and inactive ABL states. Despite certain limitations in accurately reproducing the ensembles of hidden states and precisely quantifying the changes in relative populations of the active and inactive states, we showed that combing alanine sequence masking with MSA construction and shallow MSA subsampling can capture hidden inactive states that become more stable in some of the ABL mutants while avoiding any misfolded predictions. These results are particularly significant as AF2 predictions of fold-switching allosteric systems including protein kinases are strongly biased towards predictions of the thermodynamically stable ground states and typically fail to detect less stable “excited” states. Moreover, AF2 methodologies have not been trained on mutational data and are believed to have a limited applicability for prediction of the effects of single point mutations that induced significant structural changes or unfolding.^70^ A more recent analysis of AF2 methods for predicting effects of point mutations using AF-Cluster and SPEACH_AF adaptations showed that functionally relevant structural changes in the mutational models can be obtained when mutations are introduced in the entire MSA as compared to only the input sequence.^71^ According to our analysis, another reasonable approach for capturing mutational effects on shifting conformational equilibrium an inducing dramatic structural changes is to introduce random sequence masking across the entire sequence or in specific functional regions, followed by MSA construction and shallow MSA subsampling. Although AF2 predictions can capture functionally relevant conformations the generated ensembles cannot be directly compared with the thermodynamic equilibrium ensemble of conformations. Combining the proposed AF2 adaptations showing promise in describing allosteric states and ensembles with MD simulations can enable robust characterization of structure and dynamic mechanisms by marrying the accuracy of structural predictions with the details of equilibrium ensembles resulting in a more realistic and comprehensive representation of molecular events.^72^

### Network Analysis of the AF2 Conformational Ensembles for ABL Mutants: Mutational Sites Target Allosteric Centers and Induce State-Specific Allosteric Networks

Using the ensemble-averaged model of the residue interaction networks, we computed several fundamental network properties: residue betweenness (or residue centrality) and edge betweenness. Different from previous implementations of the allosteric networks, here we employed the AF2-generated conformational ensembles of the active and inactive forms obtained for ABL mutants with alanine scanning adaptation of AF2. A global network parameter, residue centrality was used to construct the distribution profile and identify key mediating centers of allosteric interaction networks that are assumed to correspond to the profile peaks (Figure 10A-C). Although there are obvious similarities between the residue centrality distributions for the ABL states, we also noticed important differences. First, all three network centrality profiles are characterized by multiple peaks that are distributed across the entire kinase domain. However, major allosteric clusters are associated with the regulatory αC-helix (residues 292-312), the αF-helix (residues 436-453), R-spine (M309, L320, F401), 400-DFG-402 motif and 424-WTAPE-428 motif that anchors the substrate binding P+1 loop to the αF-helix, providing a plausible route for signal communication to allosteric binding (Figure 10A-C). It is well known that along with αC-helix being involved in coordination of structural changes, the αF-helix is a central scaffold for assembly of the entire molecule where the R-spine and C-spine anchor all the elements important for catalysis.^26–30^ Despite dramatic structural differences between the active and inactive states, allosteric networks may utilize the αF-helix and αC-helix along with the R-spine for mediating efficient long-range communications. At the same time, the centrality in the active state and intermediate I_1_ form showed only small peaks for C-lobe residues (Figure 10A,B) while C-lobe regions become engaged in allosteric networks in the fully inactive I_2_ form (Figure 10C).

**Figure 10.**
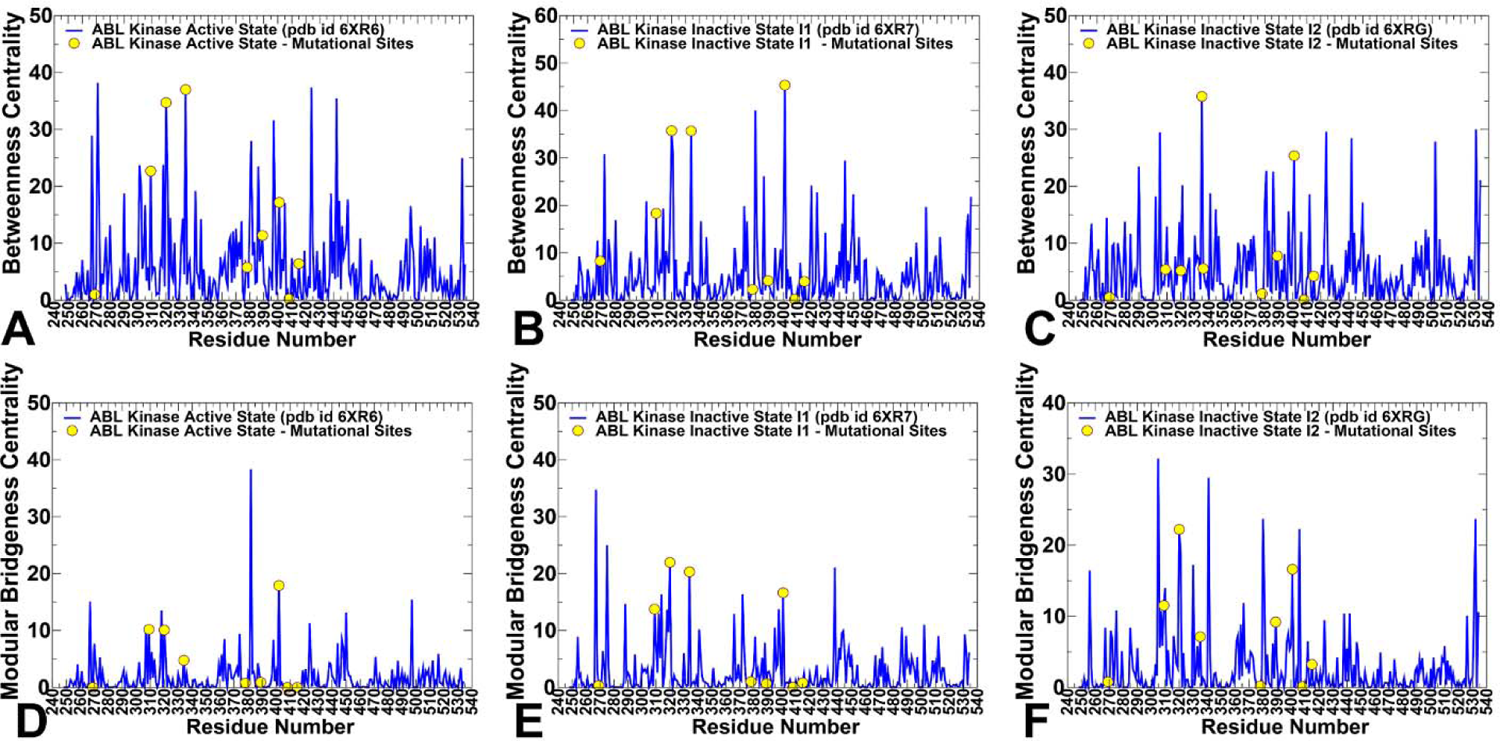
The ensemble-averaged residue betweenness centrality and bridgeness centrality profiles are obtained using conformational ensembles generated by alanine scanning AF2 adaptation for the ABL mutants. The betweenness centrality in the active ABL structure (A), inactive I_1_ state (B) and inactive I_2_ state (C). The bridgeness centrality profiles are shown in the active ABL structure (D), inactive I_1_ state (E) and inactive I_2_ state (F). The centrality profiles are shown in blue lines and positions of ABL mutations G269E, Y272H, M309L, L320I, T334I, F378, L389M, F401L, T408Y, H415P are shown in yellow-colored filled circles.

By mapping mutational sites and R-spine residues onto the centrality profiles, we noticed that M309, L320, T334 mutational positions along with F401 of the DFG motif that are fundamentally important for assembly and stabilization of the R-spine coincide with the highest centrality peaks in the active and intermediate I forms, thus corresponding to critical allosteric hotspots of long-range interactions (Figure 10A,B). Other mutational positions G269E in the P-loop as well as T408Y and H415P in the open A-loop have low centrality values and are not mediating sites of allosteric networks. These positions are located in flexible regions and can be dynamically coupled with the major allosteric centers in the global interaction network connecting the αC-helix-in, the R-spine and the substrate binding site in the active state. In network terms, the emergence of various high centrality sites that are spatially distributed typically implies a broad allosteric network with distinct communication paths connecting functional regions. A strong allosteric network dependency on these mediating sites may explain the sensitivity of the state equilibrium to M309L and L320I mutations in the R-spine leading to shift from the active to fully closed inactive structure. A noticeably different distribution of the mapped mutational position is seen in the in the I_2_ state where allosteric contributions of M309L, L320I become depleted due to disassembly of the R-spine and only T334I and F401 sites remained dominant centrality peaks (Figure 10C). In contrast to the active ABL form, we observed other mediating centers corresponding to residues Y408, Y412, L448, I451, T453, Y454 as well as C-spine residues C388, L389, and I451. The NMR analysis showed a significant role of Y408 in increasing population of the I_2_ state, and the interactions of the displaced Y412 A-loop residue with the L403/L406/M407 positions are important in allosteric changes of the inactive I_2_ form.

Using a community decomposition method the residue interaction networks were divided into local modules in which residue nodes are strongly interconnected through both dynamic and coevolutionary correlations, whereas residues that belong to different communities may be only weakly coupled. To characterize global bridges from a community structure, the community bridgeness metric was computed (Figure 10D-F). The network bridgeness profiles showed the increased modularity in the inactive forms manifested in the greater number of peaks as compared to the active state (Figure 10E,F). Among major network brightness peaks are M309L, L320I and F401 sites that are involved in transmitting allosteric signals and connecting structural clusters in the network. A number of studied ABL mutants, including P-loop mutations G269E, Y272H, A-loop mutation H415P and gate-keeper T334I are recognized as important Imatinib-resistance mutations even though some of these positions are far away from the inhibitor. NMR studies recognized that these mutations would compromise the conformational state to which Imatinib selectively binds giving rise to drug resistance.^40^ Analysis of the interaction networks has shown that residue centrality can provide insight into allosteric role of drug resistance mutations. Of special interest, the emergence T334I as dominant high centrality position in the network distributions (Figures 10,11). From the network-centric perspective, mutations in high centrality positions M309L, L320I, T334I could perturb R-spine assembly, affect the long-range interactions and alter global interaction network which may explain the state-switching impact of these modifications. We suggest that sensitivity of the ABL interaction networks to targeted perturbations of these highly connected hot spots may explain why T315I mutation and P-loop mutations could leverage a local perturbation to switch the conformational equilibrium between the inactive and active forms.

**Figure 11.**
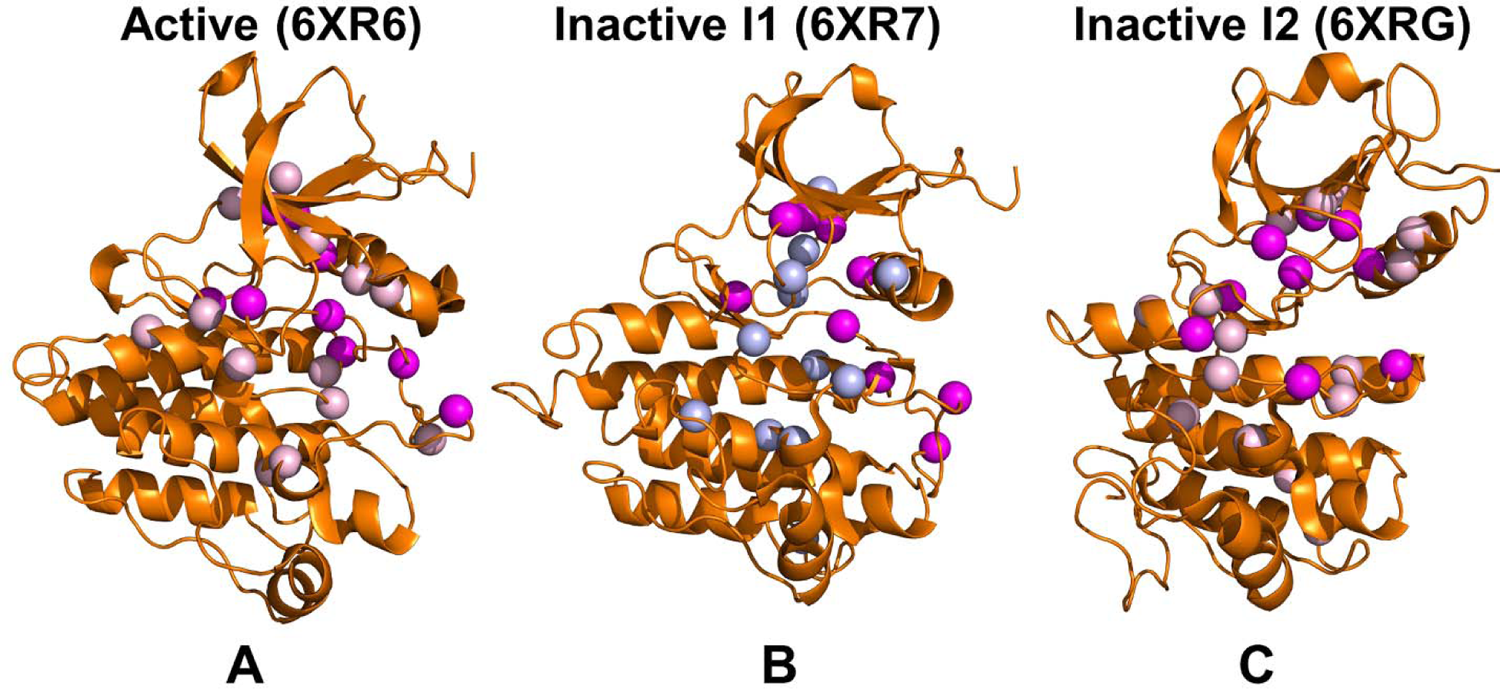
Structural mapping of network-mediating allosteric centers in the active ABL (A), inactive I_1_ state (B) and inactive I_2_ state (C). The kinase domain is shown in orange ribbons and allosteric centers are in cyan-colored spheres. ABL mutations G269E, Y272H, M309L, L320I, T334I, F378, L389M, F401L, T408Y, H415P are shown in magenta spheres.

Structural mapping of allosteric centers and sites of ABL mutations showed that mutational sites are either directly correspond to the allosteric hotspots or in local structural proximity from these positions, thus affecting the dynamics and allosteric interactions mediated by these hotspots (Figure 11). The allosteric networks are more broadly distributed in the active form (Figure 11A) and intermediate state (Figure 11B) connecting the R-spine with the highly dynamic A-loop and substrate binding site. Interestingly, structural rearrangements observed in the inactive I_2_ form produce several local clusters of allosteric centers that also include all mutational positions (Figure 11C). As a result, a more localized and narrow interaction network may be a signature of the closed inactive I_2_ state (Figure 11C) that engages a significant fraction of the A-loop residues in allosteric clusters and becomes highly dependent on all mutational sites that are allosterically coupled and act as switches of the allosteric states.

In this context, the MSAs input information in the AF2 and attention mechanism networks focused on discerning coevolving patterns are not necessarily attuned to allosteric potential and long-range effects of sequence residues. The network analysis suggests that manipulation of attention mechanism to allow for recognizing long-range dependencies between sequence positions may be helpful to detect mutation-induced structural transformations.

## Conclusions

In the current study, we examined several AF2 adaptations to characterize conformational ensembles and populations for a series of functional double and triple state-switching ABL mutants. The results of our analysis showed that widely used shallow MSA subsampling can characterize conformational heterogeneity of the ABL active form but generally is not robust for predicting mutation-induced structural changes resulting in the inactive ABL states. This study demonstrated that randomized and targeted alanine sequence masking scanning combined with shallow MSA subsampling can significantly expand the conformational diversity of the predicted structural ensembles and predict populations of both active and inactive ABL states. Consistent with the NMR experiments, the predicted conformational ensembles for M309L/L320I and M309L/H415P mutants featured the increased population of the I_2_ state. Furthermore, we found that the AF2 conformational ensembles for the G269E/M309L/T334I and M309L/L320I/T334I triple ABL mutants that share activating T334I gate-keeper substitution are dominated by the active ABL form, while G269E/M309L/T408Y mutant sampled both active and inactive conformations. The results also highlight the inherent difficulties in accurately reproducing the mutation-induced relative populations of the ABL states, as AF2 predictions are intrinsically biased towards the ground ABL state. To facilitate physics-based analysis and interpretation of the AF2 predictions, we also perform a network-based allosteric analysis of the ABL structures and AF2-generated ensembles. Through this analysis we infer that functional ABL mutations can directly target or be structurally proximal to allosteric hotpots of long-range interactions. These results underscore that learning specific patterns of coevolutionary signals together with allosteric-centric structural information on long-range couplings and attention-based learning of allosteric couplings across homologous folds may augment the predictive abilities of AF2-based methods. Combining the proposed AF2 adaptations showing promise in describing allosteric states and ensembles with MD simulations can enable robust characterization of structure and dynamic mechanisms by marrying the accuracy of structural predictions with the details of equilibrium ensembles resulting in a more realistic representation of molecular events.

## Data Availability Statement

Data is fully contained within the article and Supplementary Information material. Crystal structures were obtained and downloaded from the Protein Data Bank (http://www.rcsb.org). The rendering of protein structures was done with UCSF ChimeraX package (https://www.rbvi.ucsf.edu/chimerax/) and Pymol (https://pymol.org/2/). The software tools used in this study are freely available at GitHub sites: https://github.com/deepmind/alphafold; https://github.com/sokrypton/ColabFold/; https://github.com/RSvan/SPEACH_AF; https://www.github.com/HWaymentSteele/AFCluster; https://github.com/smu-tao-group/protein-VAE. All the data obtained in this work, the software tools, and the in-house scripts are freely available at ZENODO general-purpose open repository: https://zenodo.org/records/11204773

## Supplementary Information

Figure S1 shows structures of the ABL kinase domain in the active and inactive forms with the regulatory spine residues and mutational sites highlighted. Figure S2 describes the statistical analysis of the AF2 shallow MSA subsampling experiments and sequence coverage for MSAs for the ABL mutants. Figure S3 depicts the residue-based pLDDT profiles of the top five ranked models from the AF2 experiments with shallow MSA subsampling for the ABL mutants. Figure 4 shows the distribution density of TM scores for the predicted ABL mutant conformations using AF2 shallow MSA depth approach. Figure S5 demonstrates structural alignment of the AF2-predicted conformations for M309L/L320I mutant that are close to the inactive I_2_ state. The predicted conformations are obtained using randomized alanine sequence scanning approach combined with shallow MSA subsampling in AF2.

## Author Contributions

Conceptualization, G.V.; Methodology, N.R., M.A., G.C., H.T., S.X., G.V. P.T.; Software, N.R., H.T., S.X., M.A., G.G., P.T., G.V. Validation, N.R., G.V.; Formal analysis, N.R., G.V., M.A., G.G., H.T., S.X., and P.T.; Investigation, N.R., G.V. and P.T.; Resources, N.R., G.V., M.A, H.T., S.X., and G.V. Data curation, N.R., M.A., G.C., H.T., S.X., G.V. Writing—original draft preparation, N.R., G.V.; Writing—review and editing, N.R., G.V.; Visualization, N.R., G.V. Supervision G.V. Project administration, P.T., G.V.; Funding acquisition, P.T. and G.V. All authors have read and agreed to the published version of the manuscript.

## Acknowledgments

G.V acknowledges support from Schmid College of Science and Technology at Chapman University for providing computing resources at the Keck Center for Science and Engineering at Chapman University.

