## Supplemental Figures S1-S5 for "Prediction of Conformational Ensembles and Structural Effects of State-Switching Allosteric Mutants in the Protein Kinases Using Comparative Analysis of AlphaFold2 Adaptations with Sequence Masking and Shallow Subsampling"

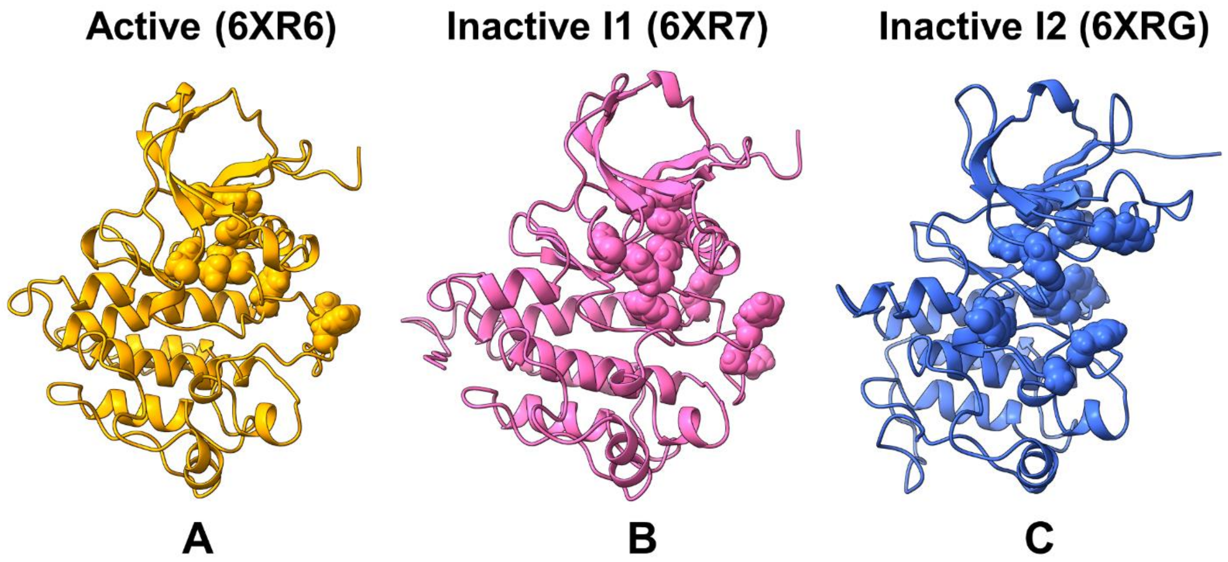


**Figure S1.** The thermodynamically stable fully active ground state of the ABL kinase domain (orange ribbons, pdb id 6XR6) (A), the inactive state $I_1$ (pink ribbons pdb id 6XR7) (B) and the closed inactive state $I_2$ (blue ribbons, pdb id 6XRG) (C). The ABL mutational sites G269E, Y272H, M309L, L320I, T334I, F378, L389M, F401L, T408Y, H415P are shown in spheres. The R-spine residues M309, L320, H380, F401 are also shown in spheres.

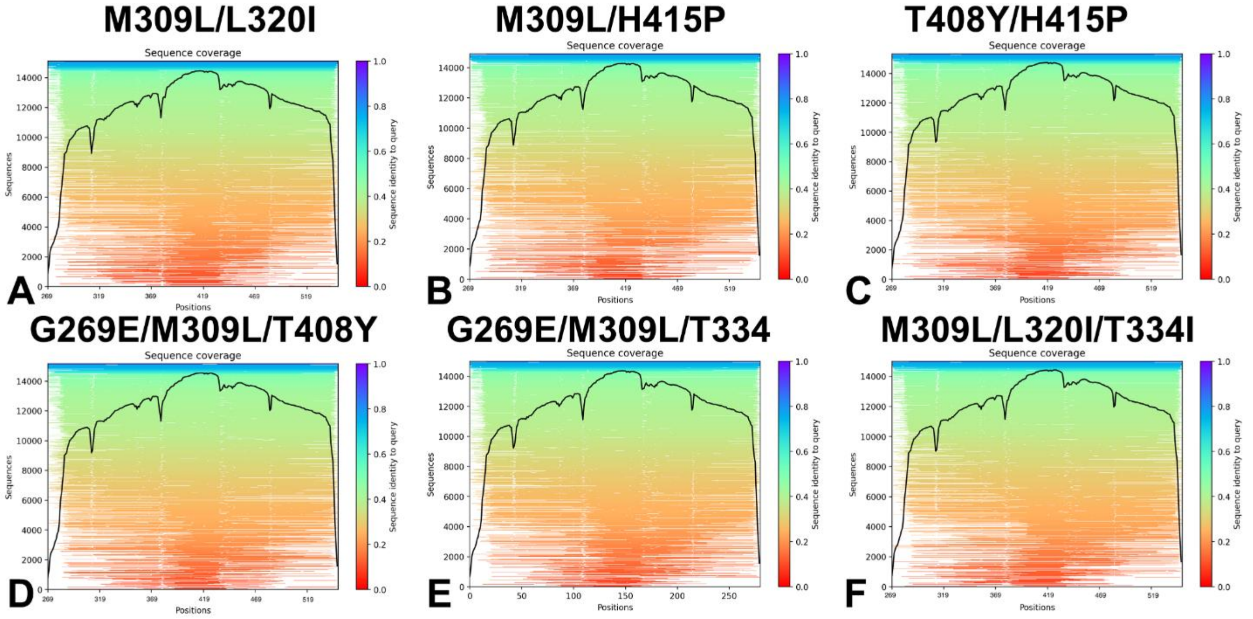


**Figure S2.** The statistical analysis of the AF2 experiments with shallow MSA subsampling for the ABL mutants : M309L/L320I mutant (A), M309L/H415P mutant (B), H415P/T408Y mutant (C), G269E/M309L/T408Y mutant (D), G269E/M309L/H415P mutant (E) and M309L/L320I/T334I mutant. A heatmap representation of the MSA indicates all sequences mapped to the input sequences. The color scale points to the identity score, and sequences are ordered from top (largest identity) to bottom (lowest identity). White regions are not covered, which occurs with sub-sequence entries in the database. The black line qualifies the relative coverage of the sequence with respect to the total number of aligned sequences.

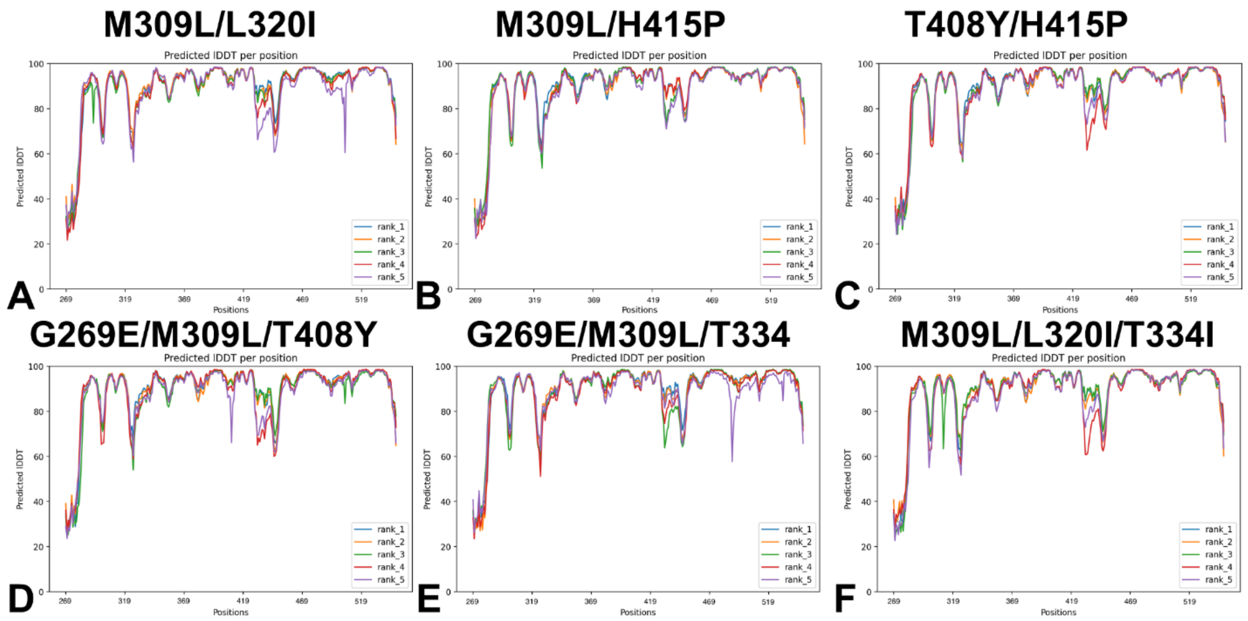


**Figure S3.** The residue-based pLDDT profiles of the top five ranked models from the AF2 experiments with shallow MSA subsampling for the ABL mutants : M309L/L320I mutant (A), M309L/H415P mutant (B), H415P/T408Y mutant (C), G269E/M309L/T408Y mutant (D), G269E/M309L/H415P mutant (E) and M309L/L320I/T334I mutant. The pLDDT and PAE profiles highlight variability for the A-loop (residues 398-421) and other functional regions.

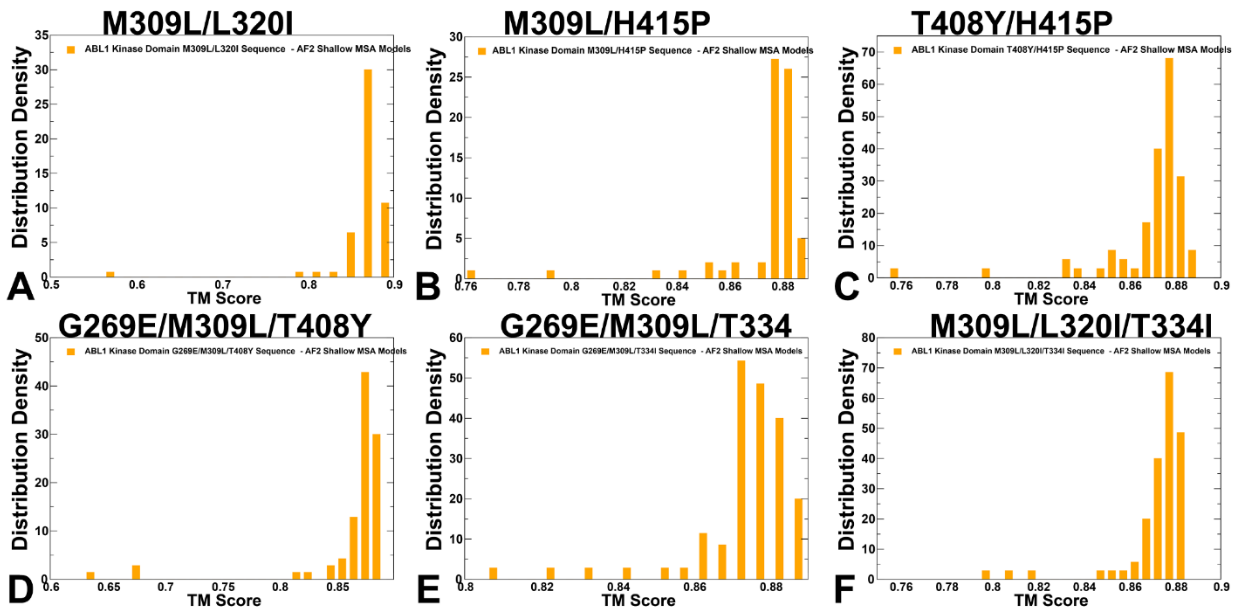


**Figure S4.** The distribution density of TM scores for the predicted ABL mutant conformations with respect to the experimental active ABL state (orange filled bars) using AF2 shallow MSA depth approach. The distributions are shown for : M309L/L320I mutant (A), M309L/H415P mutant (B), H415P/T408Y mutant (C), G269E/M309L/T408Y mutant (D), G269E/M309L/H415P mutant (E) and M309L/L320I/T334I mutant.

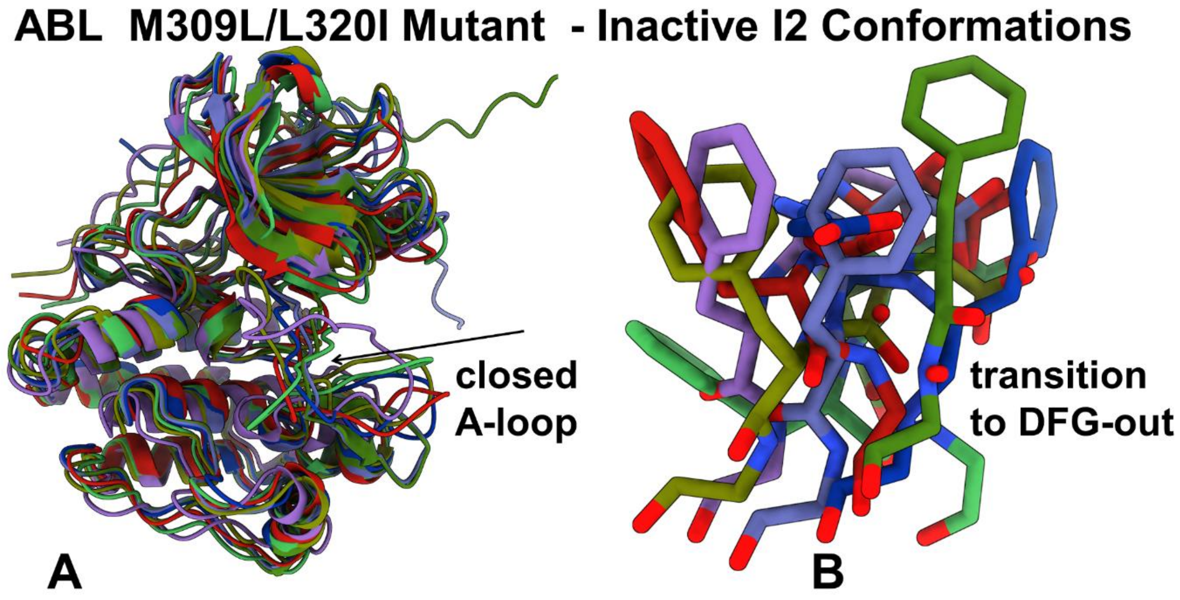


**Figure S5.** Structural alignment of the AF2-predicted conformations for M309L/L320I mutant that are close to the inactive $I_2$ state (RMSD < 1.5 Å). (A) The predicted conformations are obtained using randomized alanine sequence scanning approach combined with shallow MSA subsampling in AF2. The A-loop conformation adopts a full closed conformation as in the inactive $I_2$ experimental structure. (B) Structural alignment of the DFG motifs that adopt a number of intermediate DFG-out positions in the predicted inactive conformations.
